# A new type of transcriptional reprogramming by an IRF4 mutation in lymphoma

**DOI:** 10.1101/2022.12.29.522203

**Authors:** Nikolai Schleussner, Pierre Cauchy, Vedran Franke, Maciej Giefing, Oriol Fornes, Naveen Vankadari, Salam Assi, Mariantonia Costanza, Marc A. Weniger, Altuna Akalin, Ioannis Anagnostopoulos, Thomas Bukur, Marco G. Casarotto, Frederik Damm, Oliver Daumke, Benjamin Edginton-White, J. Christof M. Gebhardt, Michael Grau, Stephan Grunwald, Martin-Leo Hansmann, Sylvia Hartmann, Lionel Huber, Eva Kärgel, Simone Lusatis, Daniel Noerenberg, Nadine Obier, Ulrich Pannicke, Anja Pfaus, Anja Reisser, Andreas Rosenwald, Klaus Schwarz, Srinivasan Sundararaj, Andre Weilemann, Wiebke Winkler, Wendan Xu, Georg Lenz, Klaus Rajewsky, Wyeth W. Wasserman, Peter N. Cockerill, Claus Scheidereit, Reiner Siebert, Ralf Küppers, Rudolf Grosschedl, Martin Janz, Constanze Bonifer, Stephan Mathas

## Abstract

Disease-causing mutations in genes encoding transcription factors (TFs) are a recurrent finding in hematopoietic malignancies and might involve key regulators of lineage adherence and cellular differentiation^1–3^. Such mutations can affect TF-interactions with their cognate DNA-binding motifs^4, 5^. Whether and how TF-mutations impact upon the nature of binding to TF composite elements (CE) and influence their interaction with other TFs is unclear. Here, we report a new mechanism of TF alteration in human lymphomas with perturbed B cell identity. It is caused by a recurrent somatic missense mutation c.295T>C (p.Cys99Arg; p.C99R) targeting the center of the DNA-binding domain of Interferon Regulatory Factor 4 (IRF4), a key TF in immune cell-differentiation and -activation^6, 7^. IRF4-C99R fundamentally alters IRF4 DNA-binding, with loss-of-binding to canonical IRF motifs and neomorphic gain-of-binding to canonical and non-canonical IRF composite elements (CEs). Furthermore, IRF4-C99R thoroughly modifies IRF4 function, by blocking IRF4-dependent plasma cell induction, and up-regulating disease-specific genes in a non-canonical Activator Protein-1 (AP-1)-IRF-CE (AICE)-dependent manner. Our data explain how a single arginine mutation creates a complex switch of TF specificity and gene regulation. These data open the possibility of designing specific inhibitors to block the neomorphic, disease-causing DNA-binding activities of a mutant transcription factor.

## MAIN TEXT

Deregulated transcription factor (TF) activities are major contributors towards malignant transformation, and particularly promote hematopoietic malignancies. One inherent feature of such disturbed activities is the deregulation of cellular processes such as lineage adhearance, differentiation, growth or survival, thus promoting malignant transformation^1, 8, 9^. Mutations targeting TF DNA-binding motifs can affect TF:DNA interaction and/or functionality^4, 5, 10^, but whether such mutations impact upon the nature of binding to TF Composite Elements (CEs) and thus influence the interaction with other TFs is currently unclear. TF-binding to DNA is a complex process, with arginine (Arg; R)-residues playing an important role in protein-DNA recognition^11, 12^. For example, a cluster of loss-of-function *TP53* mutations affects various R-residues central to TP53:DNA interaction^13^, but also TF Interferon Regulatory Factor 4 (IRF4) contains an arginine at amino acid (AA) position 98 in the *α*3-helix of its DNA-binding domain (DBD), which is essential for its interaction with DNA^14^.

The expression of the IRF family member IRF4 is largely restricted to immune cells, where it exerts key regulatory functions^6, 7^. Due to its inherent low affinity for its cognate DNA binding motifs, IRF4 requires interaction with other TFs for efficient binding to CEs, as shown for Erythroblast transformation-specific (Ets)-IRF (EICE)^15^ or Activator Protein 1 (AP-1)-IRF CEs (AICE)^16–18^. The extent and nature of these interactions define specificity and strength of IRF4-directed transcriptional regulation in a given cell type^19, 20^. High-level IRF4 expression is characteristic for the Hodgkin/Reed-Sternberg (HRS) tumor cells of classic Hodgkin lymphoma (cHL), a common human B cell-derived malignancy^21^. However, HRS cells lack expression of the IRF4-instructed terminal B-cell differentiation program including plasma cell genes^22, 23^, and instead up-regulate genes specific to other cellular lineages^22, 24, 25^. Furthermore, HRS cells are surrounded by an inflammatory cellular infiltrate attracted by abundantly produced cytokines, chemokines and cell surface receptors^22^. Only very few genetic events driving these features of Hodgkin lymphoma cells are known.

### IRF4-C99R is recurrent in human lymphoma

By mining and integration both our own and additional published genomic and transcriptional data from well-characterized cHL cell lines, we identified and verified the same c.295T>C (chr6:394,899 T>C; hg38) variant in the *IRF4* gene in 2 of 7 HL cell lines, namely the B cell-derived HRS cell lines L428 and U-HO1 (Fig. 1a and Extended Data Fig. 1a). Based on various *in silico* analyses integrated in ANNOVAR (including SIFT, Polyphen2, MutationTaster, FATHMM, CADD score), this variant was uniformly predicted to be deleterious (Extended Data Table 1) and is completely absent in germline genomic databases (gnomAD, accessed 2022/06/16). Furthermore, no germline non-synonymous single nucleotide variants were collated in gnomAD affecting the neighboring AAs 90-104 with the exception of a singleton allele (1/251478) carrying a missense mutation in AA100. In the HL cell lines, the c.295T>C mutant allele was accompanied by at least one wild-type (WT) copy of the *IRF4* gene, and both WT and mutant *IRF4* mRNA transcripts were equally detected (Fig. 1a and Extended Data Fig. 1a). Since HRS cells are rare in the affected lymph nodes, we validated the presence of *IRF4* c.295T>C in 4 of 20 primary cHL samples representing 3 of 19 cases (16%) by DNA-PCR of laser-microdissected HRS cells (Extended Data Table 2). The S104T mutation identified in L428 cells (Extended Data Fig. 1a) was not found in the primary cases, and thus not considered as recurrent. *IRF4* c.295T>C has recently been described in Primary Mediastinal B Cell Lymphoma (PMBCL)^26^, a lymphoma entity that shares distinct biological features with cHL. Parallel mining of targeted gene panel sequencing data from an unrelated large cohort of 486 PMBCL cases identified the same *IRF4* c.295T>C mutation in 29 of the 486 cases (5.9%) (Extended Data Fig. 1b). In contrast, *IRF4* c.295T>C is not or only rarely documented in other lymphoma types (Extended Data Fig. 1b).

**Fig. 1.**
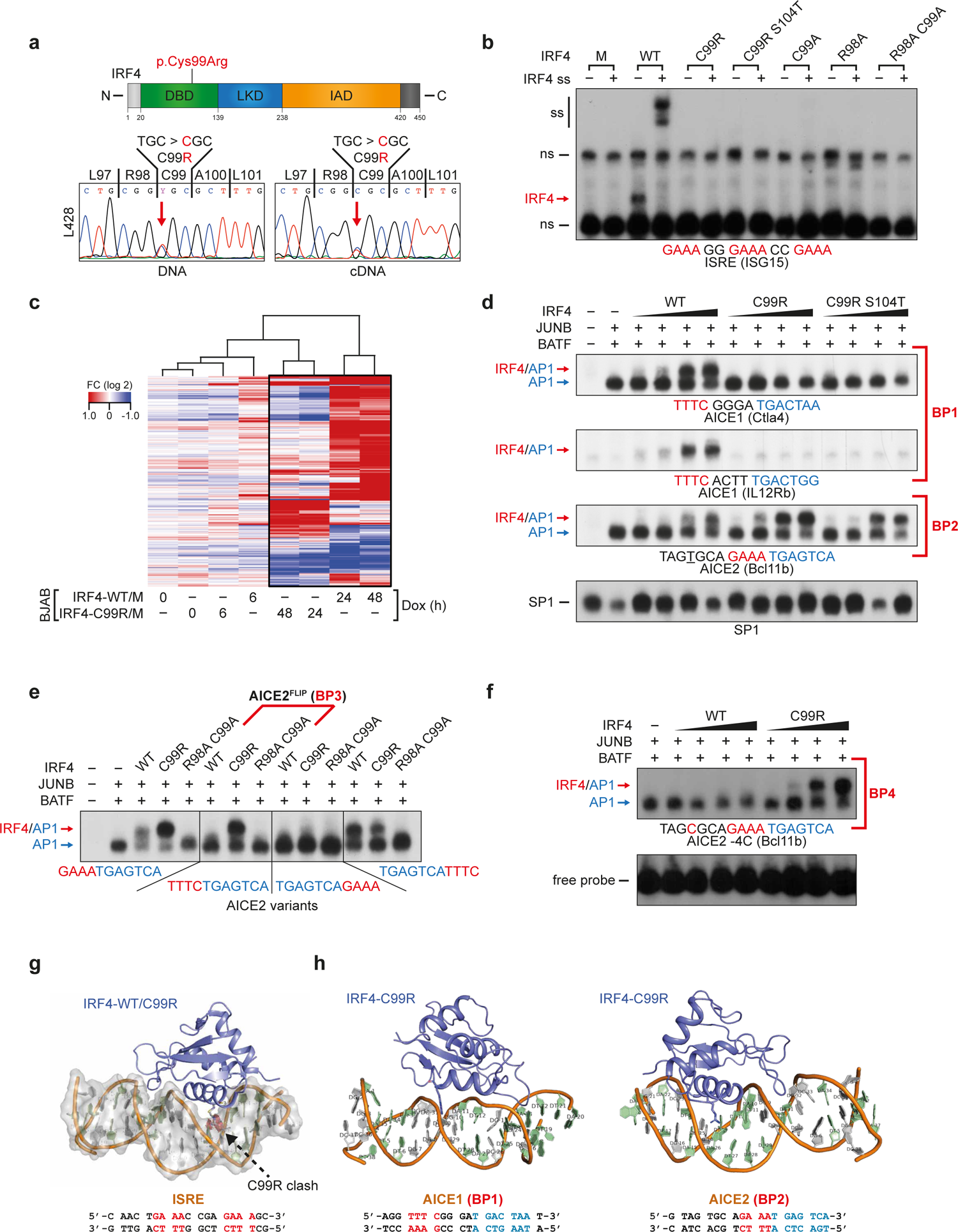
Characterization of IRF4-C99R functionality and fundamental DNA-binding alterations. (**a**) Top, scheme of IRF4 with indication of p.Cys99Arg within the DBD. Bottom, *IRF4* Sanger sequencing reads of L428 DNA (left) and cDNA (right). DBD, DNA binding domain; LD, linker domain; IAD, IRF association domain. (**b**) EMSA using the indicated ISRE probe and nuclear extracts of HEK293 cells transfected with IRF4-WT or mutants thereof. M, Mock control; C99A, R98A, and R98A C99A are known loss-of-function mutants. Red arrow, position of the IRF4-DNA complex. ss, supershift. ns, nonspecific. (**c**) Tet-inducible BJAB cells were analyzed following induction of the respective IRF4-variants for the indicated times for gene expression changes. Hierarchical clustering of 348 differentially expressed genes that change expression in either IRF4-WT or IRF4-C99R vs. Mock (M) cells. M, Mock control; Dox, doxycycline. Log2 fold changes (FC) of at least two-fold are indicated in the heat map. (**d**) Nuclear extracts of HEK293 cells transfected with JUNB and BATF and increasing amounts of IRF4 variants, as indicated, were analyzed by EMSA for binding at AICE1 (AICE1 (Ctla4) and AICE1 (IL12Rb)) and AICE2 (AICE (Bcl11b)) probes. Note the loss of IRF4-C99R and – C99RS104T binding at AICE1, marked as ‘binding pattern 1’ (BP1), and their increased binding at AICE2 (BP2). Red and blue arrows mark positions of IRF4-JUNB/BATF-DNA and JUNB/BATF-DNA complexes, respectively. EMSA of SP1 is shown as a control. (**e**) EMSAs using wild-type AICE2 (BCL11b) probe (left) or variants thereof with reverse complement IRF motif (center left, AICE2^FLIP^), or IRF motifs positioned 3’ relative the AP-1 motif (center right and right). Note the exclusive binding of IRF4-C99R to AICE2^FLIP^ (center left; BP3). Extracts and complex positions are as in (d). (**f**) Binding of JUNB/BATF together with increasing amounts of IRF4-WT or IRF4-C99R to AICE2 (Bcl11b) with a T>C mutant thereof at position −4 relative to the IRF motif. Extracts and complex positions are as in (d). All data are representative of at least three independent experiments. (**g**) Computational modelling of IRF4-C99R docking to ISRE shows that the interaction of the R99 residue with native ISRE DNA is impeded due to insufficient space in the major groove of the DNA to accommodate the long R99 side chain, as indicated by the R99 clash. (**h**) Reference free DNA modeling and docking studies with IRF4-C99R at AICE1 and AICE2. AICE1- and AICE2-DNA structure modelling shows reduced available space in the AICE1-DNA major groove when compared to AICE2, resulting in relatively poor interaction of IRF4-C99R with AICE1 DNA. This is notable for the non-canonical mode of the IRF4-C99R interaction with AICE1-DNA, where the major *α*-helix is docked onto, but not intercalated into the grooves of DNA in contrast with the AICE2-DNA interaction. This demonstrates why AICE2 but not AICE1 can interact with IRF4-C99R in EMSA analyses.

IRF4 governs at the stage of terminal B-cell differentiation the plasma cell gene expression program^27^, which largely lacks in HRS cells^23^ despite high-level IRF4 expression across all subtypes (Extended Data Fig. 1c-d). In the *IRF4* c.295T>C mutation, the basic AA arginine replaces at position AA 99 the neutral AA cysteine (Cys; C) (p.Cys99Arg; C99R), which is highly conserved in IRF4 from humans to zebrafish and also within the DBD of most other IRF family proteins (Fig. 1a and Extended Data Fig. 1e). C99R is located in the center of the *α*3-recognition helix of the DBD of IRF4 and is positioned immediately adjacent to Arg98, which is essential for specific IRF4 DNA-binding^14^. This finding suggested that C99R might interfere with the normal formation of IRF4:DNA complexes and thus with IRF4’s transcriptional activity.

### IRF4-C99R shows loss-of-function at ISRE but is functionally active

To characterize IRF4-C99R, we first explored its DNA-binding properties to the Interferon-Stimulated Response Element (ISRE) containing three consensus motifs 5’-GAAA-3’, one of the key motifs recognized by IRFs (Fig. 1b and Extended Data Fig. 1f)^28, 29^. Unlike IRF4-WT, IRF4-C99R did not bind to the ISRE at all, as demonstrated by electrophoretic mobility shift assay (EMSA). However, the recurrent nature of IRF4-C99R mutation and high-level expression in cHL suggested that this mutation does not merely constitute a loss-of-function aberration, but might possess additional, *de novo* functions. To analyze IRF4-C99R functionality, we generated tetracycline (Tet)-inducible IRF4-C99R and IRF4-WT bulk cultures of BJAB B-cell non-Hodgkin lymphoma cells, which express endogenous IRF4 only at a low level (Extended Data Fig. 2a). Time course gene expression analyses revealed that IRF4-C99R altered the expression of a clearly distinct set of genes, although fewer, compared to IRF4-WT (Fig. 1c, Extended Data Fig. 2a-e and Extended Data Table 3). Notably, IRF4-C99R was unable to induce plasma cell-specific genes (Extended Data Fig. 2f), in agreement with its lost binding capacity to the canonical ISRE motif. IRF4-C99R rescued HRS cells as efficiently as IRF4-WT from cell death induced by small-hairpin RNA (shRNA)-mediated knock-down of endogenous IRF4 (Extended Data Fig. 2g) thus corroborating its functionality.

**Fig. 2.**
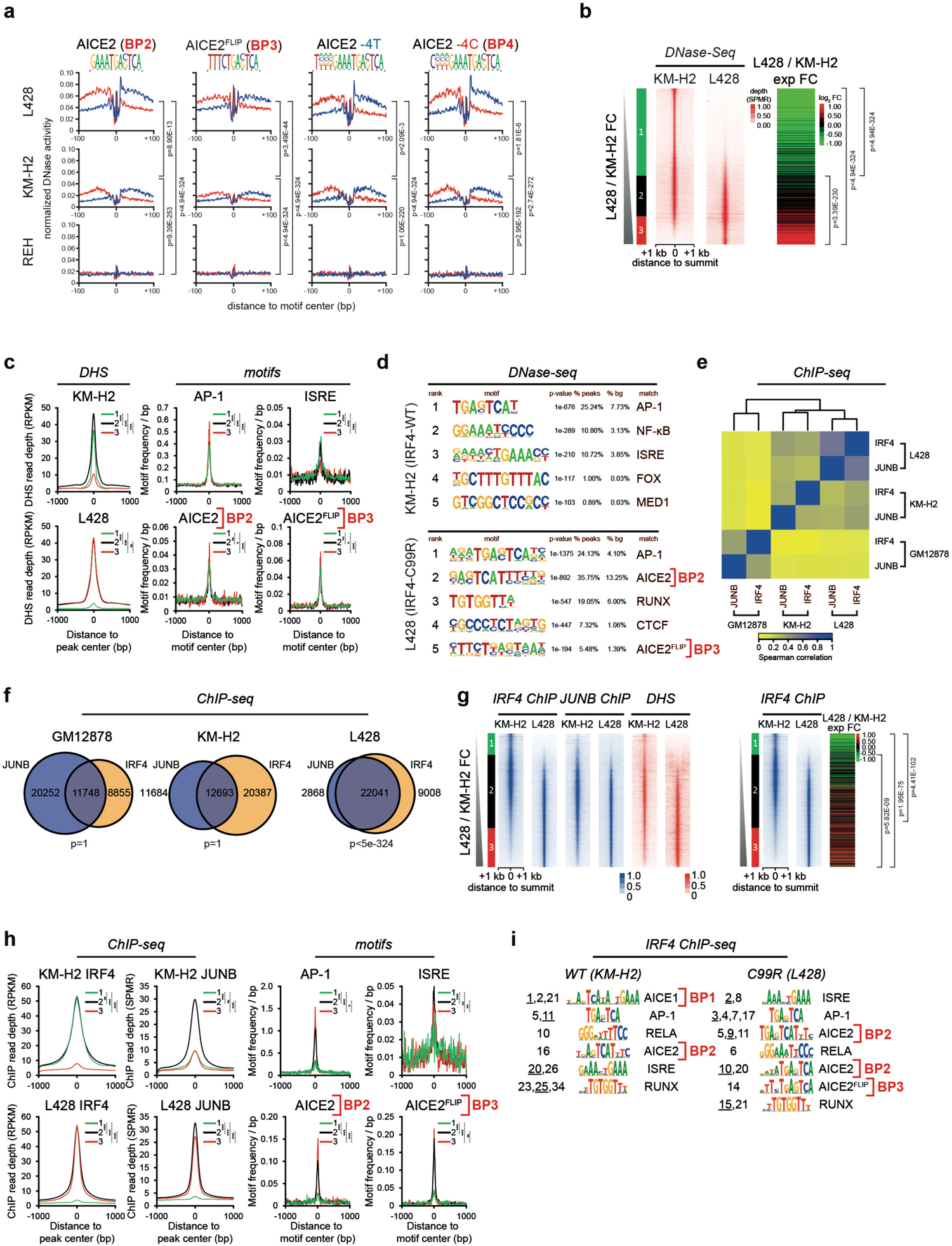
IRF4-C99R is associated with genome-wide increased and distinct DNA-binding patterns at canonical and non-canonical AICE2 sites in C99R mutation-positive lymphoma cells. (a) Digital genomic footprinting analyses showing occupancy at AICE2, AICE2^FLIP^, AICE2^-4T^ and AICE2^-4C^ sites (horizontal) in L428^IRF4-C99R^ HL cells, KM-H2^IRF4-WT^ HL and REH non-Hodgkin cells (vertical). Red and blue lines represent the forward and reverse strands. (b) Correlation between L428^IRF4-C99R^ / KM-H2^IRF4-C99R^ DNaseI-Seq fold change defining 3 classes of elements, designated groups 1 – 3, (left) and log2 gene expression fold change (right). (c) Motif frequencies in DHSs defined in (B). (D) HOMER *de novo* motif discovery in specific DHSs defined in (b). (e) Heatmap showing Spearmańs correlation clustering from IRF4 and JUNB ChIP-Seq experiments on the union of IRF4 peaks from L428^IRF4-C99R^, KM-H2^IRF4-WT^ and GM12878 cells. (f) Venn diagram-overlaps between IRF4 and JUNB ChIP peaks in GM12878, KM-H2^IRF4-WT^ and L428^IRF4-C99R^ cells. (g) L428/KM-H2 IRF4 ChIP-peak fold change analyses (left) showing corresponding JUNB ChIP peaks (center), DHSs (right) as well as gene expression fold changes (far right). (h) Motif frequencies in KM-H2^IRF4-WT^-, shared and L428^IRF4-C99R^-specific ChIP peaks. (i) *De novo* motif discovery in specific ChIP-seq datasets using ExplaiNN. Motifs are ranked by their importance (left). When more than one motif of the same class was identified, the rank of the displayed motif is underlined. Only motifs that could be annotated with a biological representation are shown. Significance levels: *, p<0.05; **, p<0.01; ***, p<0.001; ns, non significant.

### IRF4-C99R fundamentally modifies IRF4’s DNA-binding specificity

In contrast to the formation of low-affinity homodimer or multimeric complexes on ISRE DNA motifs, efficient IRF4 DNA-binding requires distinct partners such as ETS and AP-1 proteins at CEs^16, 17, 30^. Given the broad absence of ETS TFs in cHL^31, 32^, we considered the binding of IRF4 to EICE in HRS cells as being unlikely. However, constitutive AP-1 activity with high-level JUNB and BATF expression is a hallmark of HRS cells^33, 34^. We therefore speculated that IRF4-C99R regulates gene expression by DNA-binding to the recently identified AICEs, either 5’-IRF(TTTC)/nnnn/AP-1(TGASTCA)-3’ with a spacing of 4 bp (AICE1) or 5’-IRF(GAAA)/AP-1(TGASTCA)-3’ with no spacing (AICE2)^16, 35^, which both regulate key transcriptional programs in immune cells^7^. To evaluate this hypothesis, we monitored the formation of IRF4-JUNB/BATF-DNA complexes at strong (labeled as ‘AICE1 (Ctla4)’), weak (AICE1 (IL12Rb)) or intermediate (AICE2 (Bcl11b)) affinity AICE motifs^16^ (Fig. 1d and Extended Data Figs. 3a-c). While we observed a complete loss of IRF4-C99R binding at AICE1 (Ctla4) and AICE1 (IL12Rb) (designated as AICE <u>B</u>inding <u>P</u>attern 1 (BP1)), IRF4-C99R-JUNB/BATF binding at AICE2 (Bcl11b) was enhanced compared to IRF4-WT (BP2) (Fig. 1d). IRF4-C99RS104T behaved similar to IRF4-C99R (Fig. 1b and 1d, Extended Data Fig. 2g), and, as it is not recurrent, it was not included in further experiments. Strikingly, reverse complementing the IRF motif in AICE2 (Bcl11b) from 5’-GAAA-3’ to 5’-TTTC-3’ (referred to as AICE2^FLIP^) revealed an exclusively formation of mutant IRF4-C99R-JUNB/BATF-DNA complexes (Fig. 1e, AICE2^FLIP^, BP3; Extended Data Fig. 3d). Moreover, formation of AICE complexes usually requires a thymine located at −4 bp (−4T) relative to AICE2 (referred to as AICE2^-4T^)^35^. IRF4-C99R overrides this restriction, as it forms strong DNA-binding complexes in the absence of −4T, at which IRF4-WT binding is lost (Fig. 1f; AICE2^-4C^; BP4). The altered BPs of mutant IRF4-C99R were mirrored and confirmed by detailed structural modelling (Fig. 1g-h, Extended Data Fig. 4, and Extended Data Table 4).

**Fig. 3.**
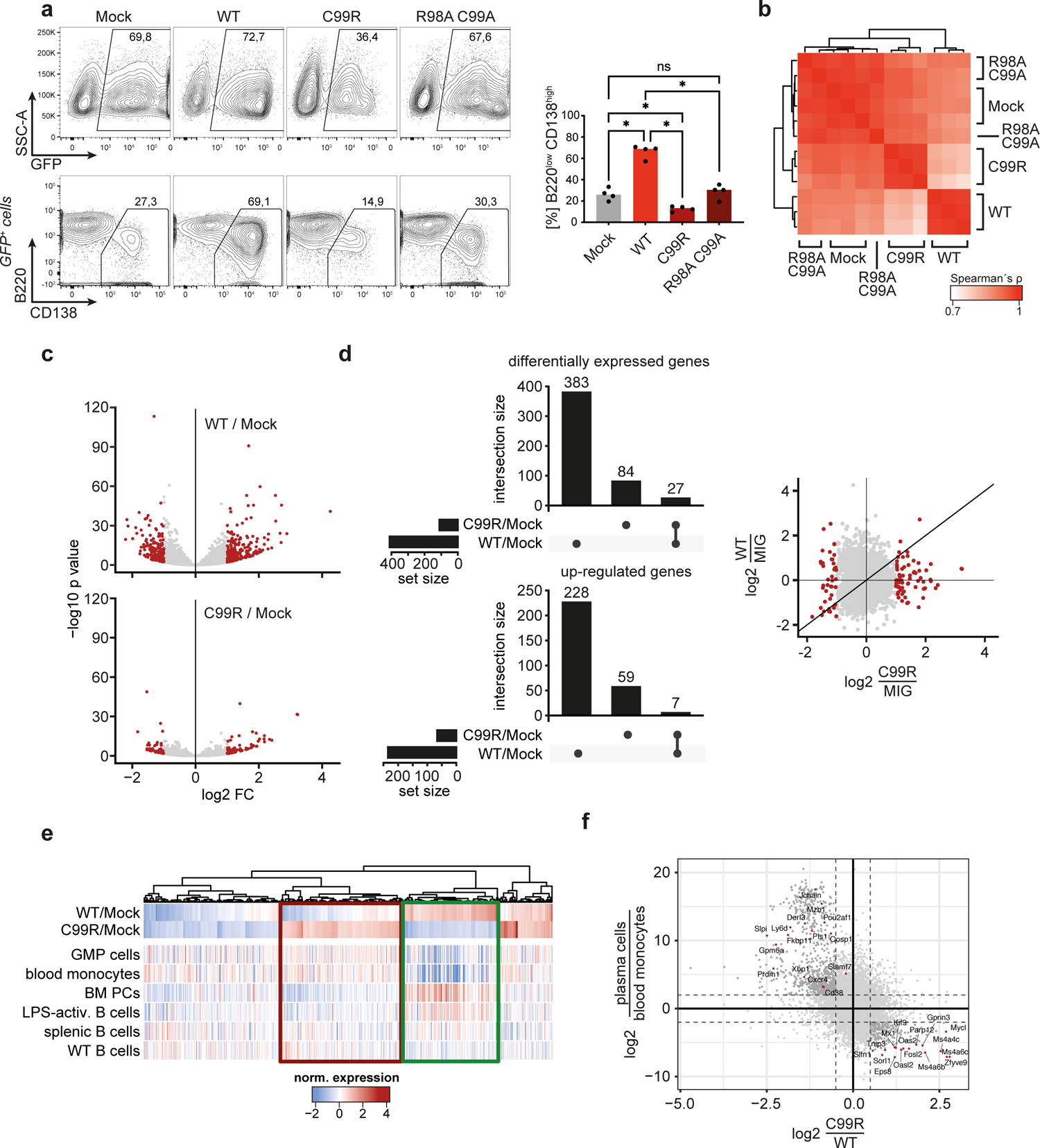
IRF4-C99R blocks IRF4-dependent plasma cell induction and regulates less but distinct genes compared to IRF4-WT. (**a**) Following culture with LPS+IL-4, C57BL/6 mouse splenic B cells were transduced with MIG control retrovirus (MIG-ctrl; Mock), IRF4-WT, IRF4-C99R or, as a further control, IRF4-R98AC99A. Transduced GFP^+^ cells were analyzed by flow cytometry for expression of CD138 and B220. Top left panels, indication of the percentage of living transduced cells in representative FACS profiles. Bottom left panels, analysis of CD138 and B220 in gated GFP^+^ cells. The percentage of CD138^high^B220^low^ cells is indicated. Right, the mean ± SD of 4 independent experiments is shown. (**b**-**f**) Isolated murine splenic B cells transduced with IRF4-WT, IRF4-C99R, IRF4-R98AC99A, or MIG-control (Mock) were analyzed by RNA-Seq. (**b**) Spearman correlation of the various samples. Note, that the Mock and the IRF4-R98AC99A-LOF transduced cells cluster together, and that IRF4-C99R clusters in between these and IRF4-WT samples. (**c**) Volcano plots of genes differentially regulated between IRF4-WT *versus* Mock and IRF4-C99R *versus* Mock (MIG-ctrl). Note, that IRF4-C99R regulates less genes compared to IRF4-WT. (**d**) IRF4-C99R and IRF-WT regulated genes show only low overlap, as shown in UpSet plots for overall differentially regulated genes (left, top panel) and up-regulated genes (left, bottom panel), as well as in overall comparisons of complete transcriptomes (right panel). (**e**) Differentially regulated genes by IRF4-C99R were compared to gene expression of lymphoid and myeloid cell types. Note, that the IRF4-C99R-dowregulated genes correspond to genes expressed in plasma cells or IRF4-WT-induced genes (green rectangle), whereas the IRF4-C99R-upregulated genes show specific expression in myeloid cells (red rectangle). GMP, granulocyte/monocyte progenitor; BM, bone marrow; PC, plasma cell. (**f**) Comparison of IRF4-C99R fold-change with ratio of gene expression from plasma cells and monocytes.

**Fig. 4.**
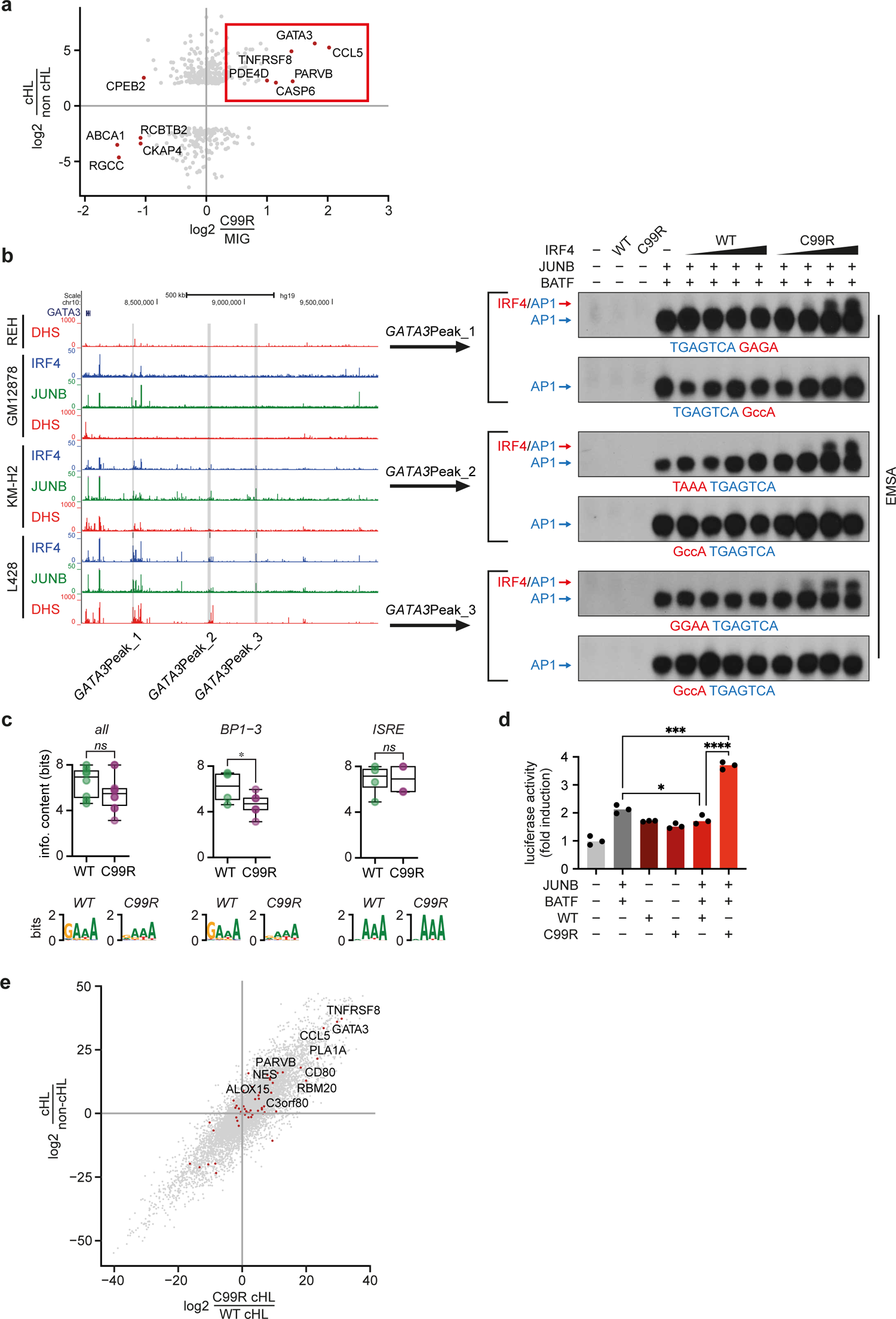
IRF4-C99R up-regulated genes encompass cHL hallmark genes in a non-canonical AICE2-dependent manner. (**a**) Comparison of fold changes between IRF4-C99R-induced genes with differentially expressed genes of Hodgkin and non-Hodgkin cell lines based on microarray gene expression analyses. Note, that IRF4-C99R induced genes include the known HL-hallmark genes *GATA3*, *CCL5* (*RANTES*) and *TNFRSF8* (*CD30*), indicated by a red rectangle. (**b**) Left, UCSC Genome Browser screenshot of REH, GM12878, KM-H2^IRF4-WT^ and L428^IRF4-C99R^ DHSs (red) as well as IRF4 (blue) and JUNB (green) ChIP peaks at the *GATA3* gene locus. L428^IRF4-C99R^-specific ChIP-peaks used for EMSA analyses are indicated by grey bars and designated as *GATA3*Peak_1, *GATA3*Peak_2 and *GATA3*Peak_3. Right, HEK293 cells were control transfected (-), or transfected with IRF4-WT, IRF4-C99R, JUNB and BATF, or combinations thereof, as indicated. Nuclear extracts were analyzed for DNA-binding activity at WT and ISRE-mutated *GATA3*Peak_1, *GATA3*Peak_2 and *GATA3*Peak_3 sites, as indicated. Note, that IRF4/JUNB/BATF complexes are only detected at the sites in the presence of IRF4-C99R, whereas IRF4-WT does not bind at these sites. (**c**) Information content (in bits; y-axis) of half-ISRE motifs within motifs identified in the ChIP-seq data of L428^IRF4-C99R^ (purple) or KM-H2^IRF4-WT^ (green) cells present in any motif (i.e. all; left), or annotated as AICE BP1-3 (center) or ISRE (right). The logos underneath of each plot represent the summary of the half-ISRE motif for each condition. (**d**) HEK293 cells were transfected with reporter construct encompassing *GATA3*Peak_1 together with AP-1 (JUNB and BATF) and IRF4 variants, as indicated. Luciferase activity is shown as fold activation compared to that of control transfected cells (far left), which is set as 1. (**e**) Comparison of gene expression changes between cell lines harboring C99R mutation (C99R / WT cHL) *versus* fold change between HL and non-Hodgkin cell lines. Note, that C99R is the primary inducer of the Hodgkin expression program in C99R bearing cells.

These patterns of alterations of the IRF4-C99R DNA-binding properties were similarly observed with recombinant proteins comprising just the DBDs of IRF4 (AA 20-139), JUNB (AA 269-329) and BATF (AA 28-87) only (Extended Data Fig. 5a-c). In addition, we visualized the DNA-bound fraction of IRF4-C99R or IRF4-WT by single-molecule fluorescence microscopy and interlaced time-lapse illumination^36^, which revealed comparable percentages of long-bound DNA contacts (> 2s) of IRF4-C99R compared to IRF4-WT molecules (Extended Data Fig. 5d). Together, our data demonstrate a unique combined loss-gain of DNA-binding preferences by IRF4-C99R, and, in particular, neomorphic binding patterns at AICE2-like motifs.

### Globally altered IRF4 DNA-binding patterns and cooperative activities in IRF4-C99R lymphoma cells

We next aimed to obtain global data supporting the above findings by interrogating our HL cell-line models. To specifically map accessible chromatin in HRS cells in detail, we first generated high-resolution genome-wide DNase I hypersensitive site (DHS) and digital footprinting data from the HRS cell lines L428, heterozygous for IRF4-C99R, and KM-H2, expressing IRF4-WT, as well as the non-Hodgkin, non-IRF4 expressing REH cells as a control (Extended Data Fig. 6a). The analyses of DNase I cutting frequencies revealed protection against DNase I digestion, indicative of occupancy by protein complexes, together with elevated accessibility of the flanking regions at AICE2 (BP2), AICE2^FLIP^ (BP3), AICE2^-4T^ and AICE2^-4C^ (BP4) only in HRS cells (Fig. 2a). Notably, and in line with our DNA-binding experiments (see Fig. 1), these motifs were highest enriched and protected in L428^IRF^^4-C99R^ (Fig. 2a). Co-localization analysis of these AICE2 motifs in L428^IRF4-C99R^ cells revealed a specific cluster corresponding to mutant-specific sites co-localizing with AP-1 motifs but not with those of other TFs typically involved in B and HL cell-gene regulation (Extended Data Fig. 6b, left), which was not observed in KM-H2^IRF4-WT^ cells (Extended Data Fig. 6b, right). These findings again support the idea of IRF4-C99R conferring cells with divergent expression profiles.

Second, to define groups of L428 or KM-H2-specific DHSs, we determined the ratio of tag counts between L428^IRF4-C99R^ and KM-H2^IRF4-WT^ cells and ranked them according to their fold change in DNaseI-seq signal (Fig. 2b; groups 1 - 3). The L428^IRF4-C99R^-specific DHSs (i.e. group 3) correlated with up-regulated gene expression in these cells (Fig. 2b). We then determined the enrichments of AICE2 (BP2), AICE2^FLIP^ (BP3), AP-1 and ISRE motifs in the different DHS groups. L428^IRF4-C99R^-specific DHSs were enriched for AICE2, AICE2^FLIP^ and AP-1 motifs, but depleted for ISRE motifs, whereas KM-H2^IRF4-WT^-specific DHSs were depleted for AICE2, AICE2^FLIP^ and AP-1 motifs, but enriched for ISRE (Fig. 2c). An unbiased search for TF motifs in the cell line-specific DHSs using HOMER revealed AICE2 and AICE2^FLIP^ as 2 of the most highly enriched motifs in L428^IRF4-C99R^-specific DHSs but not in KM-H2^IRF4-WT^-specific DHSs sites (Fig. 2d). Conversely, ISRE motifs were enriched in KM-H2^IRF4-WT^-but not L428^IRF4-C99R^-specific DHSs, again suggesting that IRF4-C99R shifts binding to distinct AICE2 motifs (Fig. 2d). Parallel DHS analyses comparing L428 *versus* REH (Extended Data Fig. 7a-b) or *versus* publicly available DHS data from lymphoblastoid GM12878 B cells (Extended Data Fig. 7c-d) also revealed specific enrichment of AICE2 motifs in L428^IRF4-C99R^-specific DHSs. Gene set enrichment analysis (GSEA) in DHSs from L428^IRF4-C99R^ *versus* KM-H2^IRF4-WT^ cells revealed an increased presence of footprinted AICE2 motif in up-regulated genes (Extended Data Fig. 7e), arguing for the functional relevance of this motif in AICE2 L428^IRF4-C99R^ cells.

A drawback of tools like HOMER is, that, although they are excellent at identifying global consensus binding motifs, they have difficulties in identifying different and slightly degenerate versions of the same motif, as found in CEs. In addition, these algorithms focus on the core motifs while ignoring the flanking nucleotides. To overcome these limitations, we used a novel deep learning tool, ExplaiNN (explainable neural networks; https://doi.org/10.1101/2022.05.20.492818), to separately discover motifs *de novo* in the cell line-specific DHS datasets. This analysis confirmed that AICE1 (BP1) in KM-H2^IRF4-WT^ and AICE2 (BP2) in L428^IRF4-C99R^ were among the most important motifs (Extended Data Fig. 8a-c).

Finally, we performed genome-wide JUNB and IRF4 Chromatin Immunoprecipitation (ChIP)-Seq analyses in L428^IRF4-C99R^ and KM-H2^IRF4-WT^ cells (Fig. 2e-i and Extended Data 9a). We included publicly available IRF4 and JUNB ChIP-Seq data from GM12878 cells, since both IRF4 and JUNB are virtually not expressed in REH cells. Sequences within IRF4-JUNB ChIP peaks clustered closely together (Fig. 2e) and showed a greater overlap (Fig. 2f) in L428^IRF4-C99R^ cells (Dice score: 0.7877) compared to KM-H2^IRF4-WT^ and GM12878 cells (Dice scores: 0.4418 and 0.4478, respectively), in line with enforced binding of IRF4-C99R to IRF and AP-1 CEs. Although IRF4-ChIP peak frequency was higher in both HRS cell lines compared to GM12878 (Fig. 2f), the overlap with JUNB was much lower in KM-H2^IRF4-WT^ cells. When individually ranked, IRF4 and JUNB showed highly similar binding patterns in L428^IRF4-C99R^ but not in KM-H2^IRF4-WT^ cells, corresponded to open chromatin regions and were associated with increased gene expression (Fig. 2g). Consistent with these analyses and motif discovery results from DHS datasets, *de novo* motif analyses by HOMER (Extended Data Fig. 9b) and supervised motif injection (Fig. 2h) showed increased frequencies of AICE2 (BP2) and AICE2^FLIP^ (BP3) motifs in L428^IRF4-C99R^-specific IRF4 ChIP peaks, while conversely showing lower ISRE motif frequencies, when compared to KM-H2^IRF4-WT^ specific ChIP peaks. These findings were also observed when IRF4 and JUNB chromatin binding patterns of L428 were compared against GM12878 cells (Extended Data Fig. 9c-d). Importantly, GSEA revealed that IRF4 and JUNB ChIP peaks were associated with increased gene expression in L428^IRF4-C99R^ but not KM-H2^IRF4-WT^ cells (Extended Data Fig. 9e).

Again, we performed *de novo* motif discovery using ExplaiNN, but this time in the ChIP-seq data sets, and found that AICE1 (BP1) was the most important motif in KM-H2^IRF4-WT^ cells, but was not identified in L428^IRF4-C99R^ cells (Fig. 2i, Extended Data Fig. 8a and Extended Data Fig. 10). AICE2 (BP2) emerged among the most important motifs in both datasets, with more importance in L428^IRF4-C99R^ cells, wherein a total of five motif types (*vs* one in KM-H2^IRF4-WT^) were identified (Fig. 2i and Extended Data Fig. 10). The analyses also revealed the unique importance of AICE2^FLIP^ (BP3) in L428^IRF4-C99R^ cells. These results agree with our DNA-binding studies (Fig. 1), and further support the notion that IRF4-C99R fundamentally alters IRF4 genome-wide DNA-binding patterns in lymphoma cells and enforces cooperative binding with AP-1/JUN TFs at distinct neo-AICEs.

### IRF4-C99R disrupts IRF4 function and exerts neomorphic gene-regulatory activity in primary B cells

To further explore the functional consequences of IRF4-C99R expression in B cells, we retrovirally transduced primary mouse C57BL/6 splenic B cells with IRF4-WT, IRF4-C99R, or the loss-of-function (LOF) variant IRF4-R98AC99A as a control (Fig. 3a and Extended Data Fig. 11a). Culturing of B cells with LPS and IL-4 led to robust endogenous IRF4 expression (Extended Data Fig. 11a) and resulted in induction of around 30% plasmablasts, characterized by a CD138^high^ and B220^low^ phenotype (Fig. 3a). The same result was obtained after expression of the non-functional IRF4-R98AC99A variant (Fig. 3a). Following ectopic expression of IRF4-WT, approx. 70% of the cells converted to a plasmablast phenotype. In contrast, IRF4-C99R reduced the number of developing plasmablasts, i.e. blocked inherent plasmablast formation, arguing for a dominant-negative function of IRF4-C99R with respect to terminal B-cell differentiation (Fig. 3a). To examine alterations in gene expression, we isolated mouse C57BL/6 splenic B cells transduced with the different IRF4 variants followed by RNA-seq analyses. Overall, the data from the respective transfectants clustered separately, with IRF4-C99R showing a transcriptional profile more similar to the R98AC99A LOF variant than to IRF4-WT (Fig. 3b). IRF4-C99R regulated a reduced set of genes (Fig. 3c), encompassing a broad loss of IRF4-WT target genes expression along with a gain of novel targets (Fig. 3d and Extended Data Fig. 11b) (Extended Data Table 5). Integration of the mRNA expression profiles of the modified splenic B cells with those from various hematopoietic cell types showed an IRF4-C99R-regulated block of overall IRF4-WT-induced and plasma cell-specific gene expression (Fig. 3e), confirming that IRF4-C99R is unable to instruct the IRF4-directed plasma cell program. Concomitantly, IRF4-C99R upregulated myeloid-associated genes (Fig. 3e-f), phenocopying a central feature of cHL tumor cells^22, 24^. Together, these data confirmed the fundamental changes in IRF4-C99R-dependent gene regulation and function compared to IRF4-WT.

### IRF4-C99R governs lymphoma hallmark gene-expression by non-canonical AICEs

To directly link IRF4-C99R-regulated genes to those specifically inherent to HRS cells of cHL, we integrated our RNA-seq data from the splenic B cells with HRS cell-specific gene expression profiles (Fig. 4a and Extended Data Fig. 12a). The latter were deduced from published microarray data as well as mRNA-seq-based gene expression profiles of cHL and non-Hodgkin lymphoma cells. Among the most prominent genes with a HRS cell-specific expression up-regulated exclusively by IRF4-C99R, but not by IRF4-WT, were *GATA3*, *CCL5* (also called *RANTES*), and *TNFRSF8* (*CD30*), all three being among the most prominent cHL hallmark genes^23, 37^, together with *CD80*, *PDE4D* and *CASP6* (Fig. 4a and Extended Data Fig. 12a).

To break down the IRF4-C99R-specific induction of these genes to the molecular level, we reanalyzed our ChIP-Seq-data for IRF4-JUNB ChIP peaks specific to L428^IRF4-C99R^ cells, but not found in KM-H2^IRF4-WT^ cells within the respective gene regulatory regions. Focusing on regions regulating *GATA3* expression, we identified several AICE2-like CEs among the L428^IRF4-C99R^-specific IRF4-JUNB ChIP peaks, designated as *GATA3*Peak_1 (5’-TGAGTCA<u>GAGA</u>-3’; the IRF-part of the binding motif is underlined), *GATA3*Peak_2 (5’-<u>TAAA</u>TGAGTCA-3’) and *GATA3*Peak_3 (5’-<u>GGAA</u>TGAGTCA-3’) (Fig. 4b, left). DNA-binding studies demonstrated that IRF4-C99R forms IRF-AP-1 composite complexes at these sites, whereas IRF4-WT did not bind to these sequences (Fig. 4b, right). Of note, none of these sites contained canonical 5’-GAAA-3’ IRF motifs, but instead non-canonical degenerate variants thereof. These results pointed to increased flexibility of IRF4 R99 compared to C99 at these motifs, similar to our observation made for the AICE2 variants described in Figs. 1e-h. Increased binding capacity of IRF4-C99R to degenerate half-ISRE containing motifs was mirrored in the observation that IRF-containing motifs identified in the ChIP-seq data using ExplaiNN were more degenerated in L428^IRF4-C99R^ cells compared to KM-H2^IRF4-WT^ cells (Fig. 4c), which was most pronounced for AICE motifs (Fig. 4c, center). We confirmed IRF4-C99R-mediated transcriptional activity of the *GATA3*P_Peak1 element by the analysis of luciferase reporter constructs (Fig. 4d and Extended Data Fig. 12b). Here, IRF4-C99R specifically enhanced the luciferase activity in combination with the AP-1 TFs JUNB and BATF, whereas IRF4-WT did not. Finally, the comparison of expression profiles of HRS cell lines harboring an IRF4-C99R mutation with those lacking IRF4-C99R in relationship with cHL-specific genes showed that the cHL hallmark genes *GATA3*, *CCL5* and *TNFRSF8* were expressed at particularly high levels in the cell lines with IRF4-C99R (Fig. 4e)

Overall, we report a somatic mutation-induced fundamental shift in TF DNA-binding specificity and motif recognition, caused by a Cys-to-Arg substitution in the *α*3-recognition helix of the DNA-binding domain of IRF4. The IRF4-C99R mutation is a hallmark of human lymphoma exhibiting perturbed B cell identity, cHL and PMBCL. IRF4-C99R is widely unable to regulate canonical IRF4 target genes, including those coordinating terminal B cell differentiation^27^. Instead, it enforces altered, disease-specific gene expression by preferred binding to canonical AICE2 sites, and by compelling neomorphic binding to non-canonical AICEs. AICE motifs are thus not only key regulatory elements of cellular differentiation and activation processes in immune cells such as T cells or dendritic cells^16–18, 38^, but we show that they can non-canonically interact with mutant TFs to establish malignancy-associated gene expression.

The arginine mutation-induced shift in DNA-binding and gene regulation highlights the critical role of Arg-residues determining interactions with the DNA interface^11, 12^. Moreover, we provide a prominent example how nucleotide sequences flanking the TF core DNA-binding motif modify TF:DNA interactions. Such an Arg-induced shift in TF binding specificity to distinct CEs, which is difficult to predict with current methodologies, distinguishes IRF4-C99R from most other mutations affecting TF:DNA interactions reported previously^4,5,10, 39, 40^ and might also operate in other diseases. We suggest to name such diseases ‘<u>M</u>utation-<u>I</u>nduced <u>N</u>eomorphic <u>Tra</u>nscription factor <u>B</u>inding’ (MINTraB)-induced diseases. Disease-causing TF activities can in principle be therapeutically targeted, which has been elaborated for a few examples like MYC or NOTCH1 (ref. ^41, 42^). Thus, the data presented herein open the possibility of designing specific inhibitors to block the neomorphic, disease-causing DNA-binding of a mutant TF, without modulating the activities of its normal counterpart.

## Supporting information

IRF4_C99R_supplementary_information

IRF4_C99R_supplementary_table_3

IRF4_C99R_supplementary_table_5

## METHODS

### Cell lines, culture conditions and transfections

HRS [L428, L1236, KM-H2, L591 (EBV^+^), U-HO1 (all of B cell origin); HDLM-2, L540, L540cy (all of T cell origin)], pro-B lymphoblastic leukemia (REH), Burkitt’s lymphoma (NAMALWA, BL-60, BJAB), diffuse large B cell lymphoma (SU-DHL-4), and HEK293 cell lines were cultured as previously described^24, 43^. Cell lines were regularly tested negative for mycoplasma contamination, and their authenticity was verified by STR fingerprinting. For preparation of nuclear extracts for DNA binding studies, HEK293 cells were transfected by electroporation in OPTI-MEM I using Gene-Pulser II (Bio-Rad) with 960 μF und 0.18 kV with 10 μg pcDNA3-FLAG-JUNB, 10 μg pcDNA3-FLAG-BATF, or increasing amounts, ranging from 0.5 to 10 μg, of the respective pHEBO-IRF4 variants. For analysis of luciferase activity, HEK293 cells were transfected with 15 μg of pGL3_*GATA3*-3P_AICE_long reporter construct, together with 150 ng pRL-TKLuc as an internal control, where indicated together with 5 μg pcDNA3-FLAG-JUNB, 5 μg pcDNA3-FLAG-BATF, or 40 μg of the respective pHEBO-IRF4 variants. 48 hours after transfection, the ratio of the two luciferases was determined (Dual luciferase kit, Promega). For generation of inducible BJAB cells, cells were electroporated with 40 μg of pRTS1-IRF4-WT or -IRF4-C99R or pRTS1 control plasmid in OPTI-MEM I using Gene-Pulser II with 50 μF and 0.5 kV. Twenty-four hours after transfection, 28 μg/mL Hygromycin B (Sigma-Aldrich, Taufkirchen, Germany) were added. After 21-28 days of culture in the presence of Hygromycin B, cells were suitable for functional assays. The respective IRF4-variants were induced by addition of 100 ng/mL doxycycline (D9891; Sigma-Aldrich).

### Preparation of whole cell and nuclear extracts, immunoblotting and electrophoretic mobility shift assays (EMSA)

Preparation of whole cell and nuclear extracts as well as immunoblotting and EMSA were performed as previously described^24, 25, 43^. For EMSA analyses, we used 3 – 5 μg nuclear extracts per lane. EMSA buffer contained 10 mM HEPES, pH 7.9, 70 mM KCl, 5 mM dithiothreitol, 1 mM EDTA, 2.5 mM MgCl_2_, 4% Ficoll, 0.5 mg/mL BSA, 0.1 μg/mL poly-deoxyinosinic-deoxycytidylic acid (poly[(dI)•(dC)]). The double-stranded oligonocleotides used for EMSA are indicated in Extended Data Table 6. After annealing, oligonucleotides were end-labeled with [*α*-^32^P]dCTP with Klenow fragment. Positions of the complexes were visualized by autoradiography. Antibodies used for supershift analyses and for immunoblotting are indicated in Extended Data Table 7.

### DNA constructs

The pHEBO-IRF4-HAtag expression construct and its control pHEBO-CMV-HAtag were kindly provided by L. Pasqualucci (New York). The R98A, C99A, C99R, and S104T mutations were introduced by use of the QuikChange Multi Site-Directed Mutagenesis Kit (Stratagene) into the pHEBO-IRF4-HAtag expression construct according to the manufactureŕs recommendations and by use of primers indicated in Extended Data Table 6. For the retroviral transduction experiments of C57BL/6 splenic B cells, the coding sequences for human *IRF4* (WT, C99R, R98AC99A) were amplified from the pHEBO-constructs using the IRF4_XhoI_forw 5’-ACCTCGAGGCCACCATGAACCTGGAGGGCGGCGGCCGA – 3’ and IRF4_EcoRI_rev 5’-ACGAATTCTTAAGGCCCTGGACCCAAAGAAGCGTAATC-3’ primers and cloned in front of the IRES sequence of the MSCV-IRES-GFP (MIG) plasmid (kindly provided by F. Rosenbauer, Münster, Germany) via XhoI and EcoRI. For the pRTS1-based inducible expression constructs^44^ of the IRF4-WT and IRF4-C99R variants, *IRF4-WT* and *IRF4-C99R* were amplified using the respective pHEBO-*IRF4* expression constructs as templates. The amplified *IRF4-WT*- and *IRF4-C99R*-products were ligated *via* XbaI into pUC19-Sfi, respectively, and mobilized by SfiI digestion for cloning into pRTS-1. For the pcDNA3-FLAG-JUNB and pcDNA3-FLAG-BATF expression constructs, full length human *JUNB* and *BATF* were amplified from cDNA of the human L428 cell line, and cloned *via* BamHI and XhoI into pcDNA3-FLAG (Invitrogen). For cloning of the pGL3_*GATA3*-3P_AICE_long reporter construct encompassing *GATA3*Peak_1, DNA from My-La cells was amplified by use of primers *GATA3*_AICE_KpnI s 5’-GCGGTACCATACAGACCCTTCCAGCCAC and *GATA3*_AICE_XhoI as 5’-GCCTCGAGAACAGATGTGGGGAGTCAGA and cloned *via* KpnI and XhoI into the multiple cloning site (MCS) of pGL3 (Promega). All constructs were verified by sequencing.

### Sanger sequencing (cell lines)

Primer sequences for the validation of *IRF4* mutations IRF4-C99R and S104T identified by whole exome sequencing in cHL cell lines were designated using the Primer3 software (http://frodo.wi.mit.edu/primer3/) (Extended Data Table 6). cDNA or RT-PCR was synthesized using the Maxima First Strand cDNA Synthesis Kit (Thermo Scientific). Sanger sequencing was performed according to standard procedures.

### Laser microdissection and PCR analyses of primary HRS cells

Tissue samples used for laser microdissection were provided by the University Cancer Center Frankfurt (UCT; Germany) and by the Hematopathology Section of Christian-Albrechts-University Kiel (Germany). Written informed consent was obtained from all patients in accordance with the Declaration of Helsinki, and the study was approved by institutional review boards and local Ethics Committees (SHN-06-2018; 15-6184-BO). Pools of 10 HRS along with pools of 10 non-tumor cells and membrane sections without tissue as controls were laser-microdissected as previously described ^45^. Following digestion with proteinase K for 3 hours at 55°C and heat inactivation for 10 minutes at 95°C, a semi-nested, two-rounded PCR with exon-spanning primers was performed to amplify exon 3 of IRF4. PCR products were separated on a 1% agarose gel. Gel-purified products were sequenced on an ABI3130 (Applied Biosystems) and evaluated with SeqScape software v2.5 (Applied Biosystems). For assessment of mutations forward and reverse sequences were mandatory. Primer sequences were (always 5’ to 3’): IRF4_E3_fw 5’-TCGTGCCACTGTACTCTAGCC; IRF4_E3_rv1 5’-ATCTGGCTGCCTCTGTTAGGT; IRF4_E3_rv2 5’-AGCTAGAAAGTGATGCTCAGAATG; IRF4_E3_fw_II 5’-AGTTCCGAGAAGGCATCGAC; IRF4_E3_rv1_II 5’-ATTGGCTCCCTCAGGAACAA; IRF4_E3_rv2_II 5’-TGTACGGGTCTGAGATGTCCA. For DNA from frozen tissue sections, the primers IRF4_E3_fw and IRF4_E3_rv2 were used in the first round of PCR (product size 389 bp), and the primers IRF4_E3_fw and IRF4_E3_rv1 in the second round (product size 346 bp). For DNA from paraffin sections, primers IRF4_E3_fw_II and IRF4_E3_rv1_II were used in the first round (fragment size 160 bp), and primers IRF4_E3_fw_II and IRF4_E3_rv2_II in the second round (fragment size 129 bp). PCR conditions were 98**°**C 4 min, 40 cycles of 98**°**C 30 sec, 62**°**C 20 sec, 72**°**C 20 sec, final elongation 72**°**C 3 min.

### *IRF4* mutation analysis in PMBCL patients

To generate a custom cRNA bait library (SureSelect, Agilent Technologies) for targeted gene capture, a total of 106 genes (including *IRF4*) that have been reported to be affected by genetic aberrations in PMBCL were selected. To ensure high quality, only samples that had a coverage of 100x in ≥80% of the exonic regions were included. The median and mean sequencing coverages were 830x and 666x, respectively. Variant calling and filtering was performed as described earlier ^46^ with the following adaptations as no germline controls were included: (i) 10%_posterior_quantile >0.1; (ii) 10%_posterior_quantile(realignment) >0.1; (iii) VAF for synonymous and nonsynonymous SNVs <0.45, >0.55 and >0.95 for regions that were not affected by SCNA. Over 3000 mutations were extensively inspected for artifacts and mapping errors through visual inspection with the Integrative Genomics Viewer (IGV). A detailed description of the PMBCL patient cohort, applied sequencing workflow and corresponding bioinformatical analysis are described in Noerenberg *et al*. (manuscript submitted). This study was conducted in accordance with the Declaration of Helsinki. The protocol was approved by the ethics review committee of the Charité – Universitätsmedizin Berlin (EA2/087/16), and of every participating center.

### *IRF4* shRNA-mediated cytotoxicity assay of L428 cells

For efficient retroviral transductions, L428 cells were engineered to express a murine ecotropic receptor as previously described^47^. Additionally, the cells were also engineered to express a bacterial tetracycline repressor allowing doxycycline-inducible small hairpin RNA (shRNA) or cDNA expression. The retroviral transduction experiments, shRNA-mediated RNA interference and cytotoxicity assays were performed as described elsewhere^47–50^. In brief, to assess toxicity of an shRNA, retroviruses that co-express green fluorescent protein (GFP) were used as described^47–50^. Flow cytometry was performed two days after shRNA transduction to determine the initial GFP-positive proportion of live cells for each shRNA. Subsequently, cells were cultured with doxycyline (40 ng/mL) to induce shRNA expression, and the proportion of GFP-positive cell was measured at the indicated time points. The GFP-positive proportion at each time point was normalized to that of the negative control shRNA and further normalized to the day two fraction. The targeting sequence of *IRF4* shRNAs #1 and #2 were 5’-CCGCCATTCCTCTATTCAAGA and 5’-GTGCCATTTCTCAGGGAAGTA as described^50, 51^. As a negative control shRNA, a previously described shRNA against *MSMO1* was used^50^. Each shRNA experiment was reproduced at least two times. For the IRF4 rescue experiments, *IRF4* (NM_002460.3) single mutant *IRF4*^C99R^ and double mutant *IRF4*^C99RS104T^ cDNAs were created and the experiment was performed as previously described^50, 51^. In brief, to assess rescue effect of an *IRF4* cDNA, L428 cells were tansduced with an *IRF4*#1 or #2 shRNA, followed by retroviral ectopic expression of either an empty vector or an *IRF4* cDNA that co-expresses GFP. We compared cell growth for each overexpression relative to the growth for the empty vector which is normalized to the 100% line, and further normalized to the day two fraction. Each experiment was reproduced at least two times. Combining the four curves (both shRNAs and their replicates) for each cDNA, aggregated curves show mean viabilities (markers) ± standard errors (transparent tunnels). At day 11, we statistically compared with 100%, i.e., with our null hypothesis for zero rescue effect (one-sample one-tailed t-tests).

### Immunohistochemistry

Formalin-fixed, paraffin embedded tissue specimens from cases diagnosed as classic Hodgkin lymphoma (30 cHL mixed cellularity subtype; 30 cHL nodular sclerosis subtype; 30 cases lymphocyte-rich subtype) were retrieved from the files of the Lymphoma Reference Centre at the Institute of Pathology, University of Würzburg, Germany. From each paraffin block 2 μm sections were cut and subjected to imunohistochemical stainings. Immunostains were performed in an automatic immunostainer using program ER2 (Bond III, Leica Biosystems, Nussloch, Germany) using the manufactureŕs protocols and detection reagents. Detection of IRF4 employed the monoclonal antibody MUM1P (M725929; DAKO/Agilent, Waldbronn, Germany).

### Cloning and purification of recombinant proteins

Codon-optimized sequences encoding the DNA-binding domains of human BATF (AA 28-87) and JUNB (AA 269-329) were cloned into pMAL-C2X (NEB). The sequence encoding human IRF4 DBD (AA 19-120) was cloned into pGEX6P1 (Cytivia). IRF4 mutations were introduced by QuikChange Site-Directed Mutagenesis Kit (Agilent) according to the manufactureŕs recommendations. All constructs were verified by sequencing. Plasmids were separately transfected into BL21-DE3-Rosetta (Novagen). Proteins were expressed overnight at 18°C in TB medium (Melford) after induction with 40 mM Isopropyl-*β*-D-thiogalactopyranosid (IPTG). Cells were resuspended in 50 mM HEPES pH7.5, 300 mM NaCl, 2.5 dithiothreitol (DTT), 1 μM DNase, 200 μM Pefablock (Carl Roth) and lysed in a microfluidizer (Microfluidics). Eluates containing MBP-fusions were applied to 5 mL amylose resin (NEB) columns and extensively washed with 20 mM HEPES pH 7.5, 150 mM NaCl, 2.5 mM DTT. Proteins were eluted in the same buffer containing additionally 10 mM maltose. Eluates containing GST-IRF4 protein were applied to a 5 mM GSH sepharose (Cytivia) column and extensively washed with 20 mM HEPES pH 7.5, 150 mM NaCl, 2.5 mM DTT. Proteins were eluted in the same buffer conaining additionally 20 mM glutathione (pH 7.5) (Sigma-Aldrich). GST was removed by the addition of Presission protease in a ratio of 1:100. All proteins were separately concentrated using 10 kD cut-off Amicon Ultra-15 Centrifugal filters (Millipore) and applied to a final gelfiltration run on a Superdex 75 column (Cytivia) using 20 mM HEPES, pH 7.5, 150 mM NaCl, 2 mM DTT as running buffer. Peak fractions containing the protein of interest were concentrated and flash-frozen in small aliquots.

### DNase-seq

DNaseI-seq was essentially performed as previously described^43^ with slight modifications. Briefly, cells were washed and resuspended at 10^8^ cells/mL in ice-cold ψ buffer (11 mM KPO_4_, pH 7.4, 108 mM KCl, 22 mM NaCl, 5 mM MgCl_2_, 1 mM CaCl_2_, 1 mM dithiothreitol, 1 mM ATP). 1 Mio REH, KM-H2 or L428 cells were treated with 12 U/mL DNaseI (Worthington) for 3 mins at 22**°**C. Digestion was stopped with the addition of 200 μl lysis buffer (100 mM Tris, pH 8.0, 40 mM EDTA, 2% SDS, 200 μg/mL proteinase K) overnight at 37**°**C. DNase digestion efficiency was checked via low-voltage overnight electrophoresis (10V) on a 0.5% TAE agarose gel. Short-fragment size-selection was performed by cutting out gel bands between 100-200 bp and subsequent purification using the QiaQuick gel extraction kit (Qiagen) according to the manufactureŕs instructions. Library preparation was performed using the KAPA hyperprep kit (Roche) following manufactureŕs guidelines. Library quality was checked via qPCR using TBP, ACTB and gene desert control oligonucleotides^52^. Libraries were sequenced at 400 million reads per library in single-end mode on separate lanes using an Illumina HiSeq 2000 system according tot he manufactureŕs instructions.

### ChIP-seq

ChIP was performed as previously described (PMID 26212328) using double-crosslinking. Cells were resuspended at 3.3×10^6^ cells/mL in PBS and first crosslinked with 8.3 μl/mL DSG (Sigma) for 45 mins at room temperature, subsequently washed 4 x and crosslinked with 1% formaldehyde for 10 mins at room temperature, with both crosslinking methods entailing sustained tube rotation. Crosslinking was quenched in 0.2M glycine and cells were washed 2x. Cells were lysed in *Buffer A* (10 mM Hepes, 10 mM EDTA, 0.5 mM EGTA, 0.25% Triton ×100), then in *Buffer B* (10 mM Hepes, 200 mM NaCl, 1 mM EDTA, 0.5 mM EGTA, 0.01% Triton ×100), at 1×10^7^ cells/mL and 4**°**C with rotation for 10 mins for both stages. Nuclei were resuspended at 2×10^7^ cells/mL in 4**°**C *IP Buffer I* (25 mM Tris, 150 mM NaCl, 2 mM EDTA, 1% Triton ×100, 0.25% SDS) and sonicated in 6x 300 μL per reaction using a Picoruptor sonicator (Diagenode) at 240W with 30 cycles of 30s on, 30s off at 4**°**C. Cell debris was pelleted via 10 min 16,000 g centrifugation and diluted in *IP Buffer II* (8.33 mM Tris, 50 mM NaCl, 6.33 mM EDTA, 0.33% Triton ×100, 0.0833% SDS, 5% glycerol final concentration). 5% of chromatin was saved as input control. Immunoprecipitation was carried out overnight using Maximum Recovery tubes (Axigen) with rotation at 4**°**C in 50 μL PBS+0.02% Tween 20 with 15 μL protein G dynabeads that were washed, blocked with 0.5% BSA and conjugated with either IRF4 (sc-6059-X, Santa Cruz) or JUNB (sc-46-X, Santa Cruz) antibodies for 4 hours ar 4**°**C with rotation. Beads were subsequently washed on ice by magnetic separation using 1xPBS+0.02% Tween 20, 2x *Wash Buffer 1* (20 mM Tris, 150 mM NaCl, 2 mM EDTA, 1% Triton ×100, 0.1% SDS), 1 x *Wash Buffer 2* (20 mM Tris, 500 mM NaCl, 2 mM EDTA, 1% Triton ×100, 0.1% SDS), 1 x with *LiCL Buffer* (10 mM Tris, 250 mM LiCl, 1 mM EDTA, 0.5% NP40, 0.5% Na-deoxycholate), 2 x with *TE/NaCL Buffer* (10 mM Tris, 50 mM NaCl, 1 mM EDTA). Beads were eluted using 2 x 50 μL *Elution Buffer* (100 mM NaHCO_3_, 1% SDS) with shaking for 15 min at RT and eluates were pooled. Chromatin was reverse-crosslinked overnight at 65**°**C in *Elution Buffer* + 200 mM NaCl, followed by 100 μg/mL RNase A and 0.25 mg/mL proteinase K digestion for 1h at 37**°**C and 55**°**C, respectively. DNA was purified *via* phenol chlorofom extaction. ChIP efficiency was checked using IL3-40, CSF1R FIRE, and gene desert control oligonucleotides^52^. Library preparation was performed using the KAPA hyperprep kit (Roche) following manufactureŕs guidelines. Libraries were sequenced in single-end mode at 50 million reads per library using an Illumina HiSeq 2000 system according to the manufactureŕs instructions.

### Single-molecule fluorescence microscopy

**(A) Cloning of IRF4 plasmids for fusion proteins**. cDNAs encoding human IRF4 and IRF4-C99R were cloned into the LV-tetO-Halo Tag plasmid using EcoRI and XbaI restriction sites and One Shot Stbl3 chemically competent E. coli (Thermo Fisher Scientific, USA)^53^. Coding regions of the plasmids were verified by Sanger sequencing. (**B**) **Generation of stable cell lines**. Lentiviral transduction was used to generate HeLa cells which stably express IRF4-WT- or IRF4-C99R-HaloTag fusion proteins^53^. In brief, HEK293T cells were transiently transfected with psPAX2 (Addgene #12260), PMD2.G (Addgene #12259) and the respective pLV-tetO IRF4-HaloTag variants using JetPrime (PolyPlus). Supernatants containing viruses were harvested through a 0.45 μm filter after 48 h. HeLa cells were infected at 37**°**C and 5% CO2 for 72 h. (**C**) **Preparation of cells for imaging**. One day before imaging, cells were seeded on a heatable glass bottom dish (DelaT, Bioptechs), and 15 min prior to imaging 3 pM silicon rhodamine (SiR) HaLoTag ligand (kindly provided by K. Johnson, Heidelberg, Germany) were added according to the HaloTag staining protocol (Promega). Thereafter, cells were washed with PBS and placed for 30 min at 37**°**C and 5% CO2 in DMEM. Before imaging, cells were washed three times with PBS and imaged in 2mL OptiMEM. (**D**) **Microscope setup**. A custom-built fluorescence microscope for single-molecule fluoresence imaging was used as described^36^. (**E**) **Interlaced time-lapse illumination and data analysis**. Cells were illuminated with a highly inclined light beam^54^ using an interlaced time-lapse illumination scheme^36^. In ITM, we repeated a pattern of two consecutive images with 50 ms camera integration time followed by a dark-time of 2 s. Localization of fluorescent molecules within an image and tracking of molecules across consecutive images was performed by use of Tracklt v1.0.1^55^. Detection and tracking parameters were: threshold factoŕ 3, ‘tracking radiu’ 2, ‘min. track length’ 2, ‘gap frame’ 0, ‘min. track length before gap fram’ 0. Molecules only detected within a single image were classified as unbound, the ones detected in two consecutive images within an area of 0.35 μm^2^ as short-bound, and those tracked over at least one dark-time as long-bound. For each imaged cell, the ratio of all bound molecules (including short- and long-bound molecules) to all molecules (including long-, short- and unbound molecules) and of long-bound molecules to all molecules was calculated. Significance between IRF4-WT and IRF4-C99R was tested with an unpaired, non-parametric t-test (Mann-Withney-test) using Graphpad prism 9.0.1.

### Reference free DNA modeling and IRF4 docking studies

To model the structural basis for the interaction of IRF-WT or IRF4-C99R with different DNA elements, unbiased random docking and interaction studies were examined using HADDOCK 2.2 (ref. ^56^) and further validation was done by docking with all outcome scores tabulated in Extended Data Table 4. Initial DNA sequence models were generated using ISRE-DNA (PDB 7JM4) template based free annealing and ternary structure prediction using HNADOCKDNA program. Deprived annealing of DNA terminals was noticed due to low dG in the modeling at the 5’or 3’. All docking studies were performed in the presence of all hydrogen atoms and water molecules (at least 5 Å around the DNA) using the Mastero package program^57^. The predicted interaction results for IRF4-WT or IRF4-C99R with different DMA elements were assessed and corroborated by analysis of all individual models in each cluster using PyMol v2.5. The binding free energies were also taken into consideration in selecting the best possible models. Further validation and refinement were completed by ensuring that the residues occupied Ramachandran favoured positions using Coot (https://www2.mrc-lmb.cam.ac.uk/personal/pemsley/coot/). All structures were visualized, and the figures were generated using PyMol.

### RNA-seq of human lymphoma cell lines

RNA-seq analyses of L428, L1236, KM-H2, U-HO1, L591, HDLM-2, L540, L540cy, REH, NAMALWA, BJAB and SU-DHL-4 cells was performed in duplicates. In brief, barcoded mRNA-seq cDNA libraries were prepared from 600 ng of total RNA using Illuminás TruSeq Stranded RNA Sample Preparation Kit. mRNA was isolated using Oligo(dT) magnetic beads. Isolated mRNA was fragmented using divalent cations and heat. Fragmented mRNA was converted into cDNA using random primers and SuperScriptII (Invitrogen). This was followed by second strand synthesis. cDNA was repaired and 3’ adenylated. 3’ single T-overhang Illumina multiplex specific adapters were ligated on the cDNA fragments, and these fragments were enriched by PCR. All cleanups were done using Agencourt XP magnetic beads. Barcoded RNA-seq libraries were clustered on the cBot using the Truseq PE cluster kit V3 using 10 pM and 2 x 50 bps were sequenced on the Illumina HiSeq2500 using a Truseq SBS V3 kit.

### Generation of retroviral particles, mouse B cell isolation, retroviral transduction

By use of calcium-phosphate buffer, 10 μg of retroviral plasmids (MSCV-based) encoding human IRF4-WT, IRF4-C99R, or IRF4-R98AC99A were transfected into the Plat-E packing cell line^58^, together with packaging plasmids pGagpol (10 μg) and pEnv (2 μg) (both courtesy of A. Leutz, Berlin) and 25 μM chloroquin (#6628, Sigma). Thereafter, cells were incubated for 6-8h at 37**°**C and 5% CO2, followed by change of medium to B cell medium (DMEM high glucose (4,5 g/L) supplemented with 10% FCS, 1% sodium pyruvate, 1% penicillin-streptomycin, 1% HEPES, 1% L-Glutamin, 1% non-essential amino acids and 0.05% *β*-mercaptoethanol) and further cultivation. 48h after transfection, cell culture supernatants were harvested, filtered (0.45 μm) and frozen at −80**°**C. Splenic B cells were isolated from C75BL/6 mice by CD43 depletion with magnetic anti-mouse CD43 microbeads (#130-049-801, Milenyi Biotech) according to the manufactureŕs instructions. Purified B cells (density 1×10^6^ cells/mL; 4×10^6^ cells per well) were cultured in the presence of recombinant mouse IL-4 (25 ng/mL; #404-ML, R&D) and LPS (20 μg/mL; #L2880, Sigma) over night to induce B cell activation and terminal differentiation. 24h after isolation, B cells were collected (300xg, 5 min, 4**°**C) and resuspended in B cell medium supplemented with 8 μg/mL polybrene (#TR-1003, EMD Millipore) at a density of 2×10^6^ cells/mL. To introduce the IRF4 variants, 4×10^6^ B cells per well were plated in 2 mL on 6-well plates that had been coated with RetroNectin (25 μg/mL, 4**°**C, overnight; #T100B, Takara), blocked with 2% BSA in PBS (1h) and pre-loaded with the respective retroviral particles (1h, 37**°**C) Retroviral transduction was performed by the addition of 2 mL of the respective retroviral supernatant and subsequent centrifugation (800xg, 90 min, 32**°**C). 24h after transduction, B cells were collected (300xg, 5 min, 4**°**C), resuspended in B cell medium and cultured (density 1×10^6^ cells/mL; 4×10^6^ cells per well) for another 72h (FACS for RNA-seq, flow cytometric analysis of plasma cell differentiation) in the presence of recombinant mouse IL-4 and LPS.

### Flow cytometry of C57BL/6 splenic B cells

Retrovirally transduced B cells were harvested, blocked with TruStain FcX (*α*-mouse CD16/32; 10 min, 4**°**C; #101320, BioLegend) and stained (20 min, 4**°**C) with B220-PerCP/Cyanine5.5 (#103235; BioLegend) and CD138-PE (#142504; BioLegend) in PBS, pH7.2, supplemented with 3% FCS and 1 mM EDTA. Analysis of the samples was performed on a FACSCantoII instrument (BD BioSciences) or sorted on a FACSAria (BD BioSciences). FlowJo software (BD FlowJo, RRID:SCR_008520; v9.9.6) was used to generate plots.

### Bioinfomatics analyses of HL RNA-seq, DNase-seq and ChIP-seq data

#### HL cell line RNA-seq processing

Reads were aligned in paired-end mode to the hg19 genome using STAR v2.3.0 (ref. ^59^) using --outSAMattributes Standard --outSAMunmapped None -- outReadsUnmapped Fastx --outFilterMismatchNoverLmax 0.02 as parameters. Counts were obtained using featureCounts v2.0.0 (ref. ^60^) with -p -B -C -Q 10 --primary -s 0 as parameters. Normalisation and differential gene expression analysis was performed using DESeq2 v1.14.1 (ref. ^61^) using the standard analysis protocol, performing variance stabilisation transform normalisation. Gene set enrichment analyses were performed using GSEA v3.0 (ref. ^62^).

#### DNase-seq and ChIP-seq processing

Base-calling was carried out using HiSeq Analysis Software v2.0 (Illumina) Reads were demultiplexed using bcl2fastq v2.16.0 (Illumina). As libraries were sequenced in separate lanes, unassigned reads/unreadable indexes were assigned to their respective lane. Reads were subsequently aligned in single-end mode to the hg19 genome using bowtie2 v2.1.0 (ref. ^63^) using --very-sensitive-local as a parameter and sorted by coordinate ordering using samtools sort v1.1 (ref. ^64^). Peak calling and depth coverage track generation were carried out using macs2 v2.1.0 (ref. ^65^) using callpeak -g hs -q 0.001 -B -- SPMR --trackline <trackline> as parameters, with --keep-dup all and --keep-dup auto for DNaseI- and ChIP-Seq assays, respectively, to account for high depth sequencing of DNase-Seq libraries. Peak calling yielded 61983, 65612 and 68370 peaks for Reh, KM-H2 and L428 DNase-Seq datasets, respectively, 33082 and 30022 peaks for KM-H2 and L428 IRF4 ChIP-Seq datasets, respectively, as well as 24914 and 24886 peaks for KM-H2 and L428 JUNB ChIP-Seq datasets, respectively.

#### DNase-seq and ChIP-seq processing

For pairwise comparisons, the union of peak summits was obtained as previously described^66^ and masked against blacklisted and simple repeat regions ^67^ using bedtools intersect v2.19.0 (ref. ^68^) with -v as a parameter. Corresponding depth coverages were obtained using homer annotatePeaks v4.6 (ref. ^69^) with -hist 10 -ghist -size 2000 as parameters and subsequently log_2_ fold-changed ranked on total signal [-100 bp; +100 bp] around peak summits as previously described^70^. Heatmaps were obtained using Java TreeView v1.1.4 (ref. ^71^). Venn diagramme overlaps and specific peak populations were computed using ChIPpeakAnno makeVennDiagram v.1.12.0 (ref. ^72^) and bedtools intersect with totalTest=<sum of ChIP-Seq peak numbers> and -u as a parameter, respectively. Spearman correlation clustering heatmaps were obtained via gplots heatmap.2 v2.17.0 using total ChIP-Seq signal [-100 bp; +100 bp] around the union of GM12878, KM-H2 and L428 IRF4 ChIP-Seq peak summits. For motif, DNase-Seq and ChIP-Seq average profile significance testing, signal [-200 bp; +200 bp] from summits were averaged per region, split into three classes and tested for significance using t-tests. For GSEA analyses, peaks were annotated to the closest gene using bedtools closest using -t first as a parameter; the top 1000 specific peaks sorted by decreasing signal were used. Footprinted motif co-occurence clustering was performed as previously described^66, 70^, with specific peaks and 1,000 similarly-sized samplings of control peaks being annotated with 16 motifs using Homer annotatePeaks. Intersection matrices were computed using pyBedtools intersection_matrix^73^. Enrichment z-scores were computed by subtracting mean co-occurrences between the specific peaks and control peaks, dividing by the standard deviation of control peaks.

#### Public dataset processing

Reads for IRF4 and JUNB GM12878 ENCODE ChIP-Seq datasets^74, 75^ and GM12878 ENCODE DNase-Seq^76^ were retrieved from the Sequence Read Archive (SRA) and processed as ChIP-Seq datasets above. L428 and REH DNase-seq datasets generated in this study were complemented in reads from corresponding previously published, lower sequencing depth datasets^43^.

### Motifs discovery, average profiles and heatmaps

Motif discovery was performed using Homer findMotifsGenome using default parameters. Motif average profiles and heatmaps were generated using Homer annotatePeaks using -hist 10 -ghist -size 2000 as parameters and plotted using Java TreeView.

### Digital genomic footprinting

Digital genomic footprinting was performed using pyDNase wellington_footprints v0.2.6 (ref. ^77^) using -A as a parameter, yielding 60669, 75813 and 75755 footprints for REH, KM-H2 and L428 cells, respectively. Individual motif bed files were obtained from union of REH, KM-H2 and L428 footprints annotated for each motif using Homer annotatePeaks -mbed -size given as parameters and subsequently plotted as motif footprint profiles using pyDNase dnase_average_profiles with -A -n as parameters. AICE2 and AICE2^FLIP^ motifs were obtained from ref. ^35^. Motif footprinting scores were obtained using wellington_score_heatmap using -A as a parameter, with scores at the footprint centre being log2 transformed and used for t-test significance testing^78^.

### Bioinformatic analysis of mouse splenic B cell and lymphoma cell line RNA-seq data: RNA-seq of splenic B cells

RNA prepared from isolated murine splenic B cells transduced with MIG control virus or IRF4-WT, IRF4-C99R, or IRF4-R98AC99A variants was processed by use of the KAPA mRNA Hyper Prep Kit for Illumina Platforms (KK8580; Roche) and KAPA Single-Indexed Adapter Kit (KK8700; Roche). Libraries were sequenced by use of Illumina HighSeq 4000. RNAseq data from mouse splenic B cells was processed using PiGx-RNA-seq^79^ pipeline. In short, the data was mapped onto the GRCm38/mm10 version of the mouse transcriptome (downloaded from the ENSEMBL database^80^) using SALMON^81^. The quantified data was processed using tximport^82^, and the differential expression analysis was done using DESeq2 (ref. ^61^). Genes with less than 5 reads in all biological replicates of one condition were filtered out before the analysis. Two groups of differentially expressed genes were defined - a relaxed set containing genes with an absolute log2 fold change of 0.5, and a stringent set containing games with an absolute log2 fold change of 1. The fold change was deemed significant if the adjusted p-value was less than 0.05 (Benjamini-Hochberg corrected).

### RNA-seq of lymphoma cell lines

RNAseq data of the Hodgkin and non-Hodgkin cell lines was processed using the PiGx-RNA-seq^79^ pipeline. In short, the data was mapped onto the GRCh38/hg38 version of the human transcriptome using SALMON^81^. Differential gene expression results were integrated with the previously analyzed microarray data from cHL cell lines^83^.

### Data integration and visualization

Splenic B cell per-sample heatmap was constructed by calculating the Pearson correlation coefficient DESeq2 normalized expression values. The heatmap was visualized using the ComplexHeatmap package^84^. The number of stringently differentially expressed genes in each condition was visualized using UpSet diagrams^85^. Human and mouse genes were mapped through the orthologous assignment using the ENSEMBL database. Monocyte, B cell, and plasma cell expression profiles were extracted from the ARCHS4 database^86^. Samples with the following keywords in the Sample_source_name_ch1 field were used in the analysis: ‘granulocyte-monocyte progenitor (GMP) cells’, ‘Blood-derived monocyte’, ‘Bone marrow, plasma cells, WT’; ‘Splenic B cells, Wild type’; ‘WT, B cells’; ‘LPS activated B cells’. If multiple samples corresponded to one condition, their expression values were averaged.

### Microarray analyses of inducible BJAB cells

For microarray gene expression analyses of BJAB cells following tet-induction of IRF4-WT or IRF4-C99R, mRNA was processed by use of the Illumina Total Prep RNA Amplification Kit (AMIL1791; Invitrogen) and use of Human HAT-12_v4 Bead Chips (Affimetrix). The microarray gene expression data from a total of 24 arrays were analysed in GenomeStudio software (Illumina, Little Chesterford, UK) with background subtraction from experiments that were performed with Tet-inducible BJAB cells expressing Mock control, IRF4-WT or the IRF4-C99R mutant. The raw data output by GenomeStudio was analysed using the Lumi R package^87^ with quantile normalisation. The 10% threshold (p value <= 0.1) was applied to all samples. Genes with at least two-fold-change in expression (either up or down) were selected in either IRF4-WT *versus* Mock and IRF4-C99R *vs* Mock in different time courses. Principle component analysis and hierarchical clustering were carried out using all expressed genes across all replicate samples to show that replicates are highly correlated and then hierarchical clustering of differentially expressed genes was carried out only for genes associated with at least a two-fold change in one condition in either IRF4-WT *versus* Mock and IRF4-C99R *vs* Mock. Hierarchical clustering was used with Euclidean distance and average linkage clustering.

## ExplaiNN models and Calculation of Information content

### Deep learning models

Four different ExplaiNN models (https://doi.org/10.1101/2022.05.20.492818), each with 100 units, were trained on either IRF4-C99R or IRF4-WT ChIP-/DNase-seq data. The architecture of each unit was as follows: • 1st convolutional layer with 1 filter (26×4), batch normalization, exponential activation to improve the representation of the learnt sequence motifs ^88^ and max pooling (7×7); • 1st fully connected layer with 100 nodes, batch normalization, ReLU activation and 30% dropout; and • 2nd fully connected layer with 1 node, batch normalization and ReLU activation. For training the models, ChIP-/DNase-seq peaks were resized to 201 bp by extending their summits 100 bp in each direction using BEDTools slop (version 2.30.0)^68^. Negative sequences were obtained by dinucleotide shuffling each dataset using BiasAway (version 3.3.0)^89^. Sequences were randomly split into training (80%), validation (10%) and test (10%) sets using the “train_test_split” function from scikit-learn (version 0.24.2) (https://jmlr.csail.mit.edu/papers/v12/pedregosa11a.html). Models were trained as described in ExplaiNN. Briefly, using the Adam optimizer (https://arxiv.org/abs/1412.6980) and binary cross entropy as loss function, applying one-hot encoding, setting the learning rate to 0.003 and batch size to 100, and using an early stopping criteria to prevent overfitting. Models were also interpreted following the specifications from ExplaiNN. The filter of each unit was converted into a motif by aligning all sub-sequences activating that filter’s unit by ≥50% of its maximum activation value in correctly predicted sequences. The importance of each motif was calculated as the product of the activation of its unit for each correctly predicted sequence activating that unit by ≥50% of its maximum activation value times the weight of the final layer of that unit.

### Information content

For each ChIP-seq motif annotated as AICE1, AICE2 or AICE2^FLIP^, the information content for the four bp corresponding to the half-ISRE site was calculated using Biopython^90^. Summary motifs of the half ISRE sites in present in these motifs were obtained by aligning the individual 4-mers corresponding to the half ISRE sites. Statistical significance was computed using the Welch’s t test (one-tailed) as implemented in SciPy (version 1.7.1)^91^.

### Data availability

All data presented in this study is available at the Gene Expression Omnibus^92^ under superseries accession GSE211445. Deposited datasets were DNaseI-Seq: GSE211441; ChIP-Seq: GSE211443; HL and NHL cell line RNA-Seq: GSE211444; BJAB cells with Tet-inducible control, IRF4-WT and IRF4-C99R Illumina BeadChip HT-12 V4.0 expression arrays: GSE211913.

## ACKNOWLEDGEMENTS

The authors like to thank Natalia Soloch (Poznan), Sabine Werner (Berlin) and Brigitte Wollert-Wulf (Berlin) for excellent technical assistance, and P. Rahn (Berlin) for cell sorting. Support for infrastructure has been provided by the KinderKrebsInitiative Buchholz/Holm-Seppensen. We thank Wolfram Klapper (Kiel) for providing HL tissue samples, and Patrick Sorn (Mainz) and the Team Medical Genomics (TRON gGmbH, Mainz) for RNA-seq processing and data analysis. RS and MG received funding from the European Uniońs Horizon 2020 research and innovation programme under grant agreement No 952304. OF and WWW were supported by grants from the Canadian Institutes of Health Research (PJT-162120), Natural Sciences and Engineering Research Council of Canada (NSERC) Discovery Grant (RGPIN-2017-06824), and BC Childreńs Hospital Foundation and Research Institute. RK and M-LH were supported by the Wilhelm Sander Foundation (2018.101.1). This study was in part supported by grants from the Brigitte and Dr. Konstanze Wegener-Stiftung (#55), the Deutsche Krebshilfe (#70113148, #70113643) awarded to FD, funds of the Max Planck Society (K150) to RG, the Deutsche Forschungsgemeinschaft to MJ and SM (MA 3313/2-1 and JA 1847/2-1). Work in the lab of CB and PNC was funded by a Blood Cancer Research UK programme grant (15001), the Kay Kendall Leukemia Fund (KKL725) and a studentship donation from Arthur D. Riggs from City of Hope for BE-W.

## AUTHOR CONTRIBUTIONS

NS, PC, VF, MG, and OF designed and performed experiments, interpreted data and wrote the manuscript; NV, MGC, and SS performed and interpreted structure modelling; SA analyzed and interpreted microarray data; MAW, M-LH, SH, and RK designed, performed and interpreted HRS single cell analyses; IA and AR performed and interpreted IHC analyses; FD and DN performed and interpreted PMBCL analyses; OF and SG designed and performed production of recombinant proteins; CMG and AR designed, performed and interpreted single-molecule fluorescence microscopy; TB analyzed RNA-seq data; EK and SL performed experiments and interpreted data; UP, WW, MC performed experiments, interpreted data and contributed to writing of the MS; AP interpreted data; B-EW, LH, PC, and NO designed, perfomed and interpreted DNase and ChIP experiments; AW, WX, MG and GL designed performed and interpreted shRNA experiments; KS, KR, GL, AA, and WWW, interpreted data and contributed to writing of the manuscript; PNC, CS, RS, RK, RG, MJ and CB designed research, interpreted data and wrote the manuscript, SM designed research, interpreted data, wrote the manuscript and supervised the project. All authors discussed the results and commented on the manuscript.

## ADDITIONAL INFORMATION

This work contains Extended Data Figures 1 to 12 and Extended Data Tables 1 to 7.

## CORRESPONDING AUTHOR

Stephan Mathas, MD; Max-Delbrück-Center for Molecular Medicine and Charité – Universitätsmedizin Berlin, Hematology, Oncology and Cancer Immunology; Robert-Rössle-Str. 10; D-13125 Berlin, Germany; Tel.: +49.30.94063519; Fax: +49.30.94063124.

