## Supplementary material for "A new type of transcriptional reprogramming by an IRF4 mutation in lymphoma": IRF4_C99R_supplementary_information

#### Extended Data Figure 1

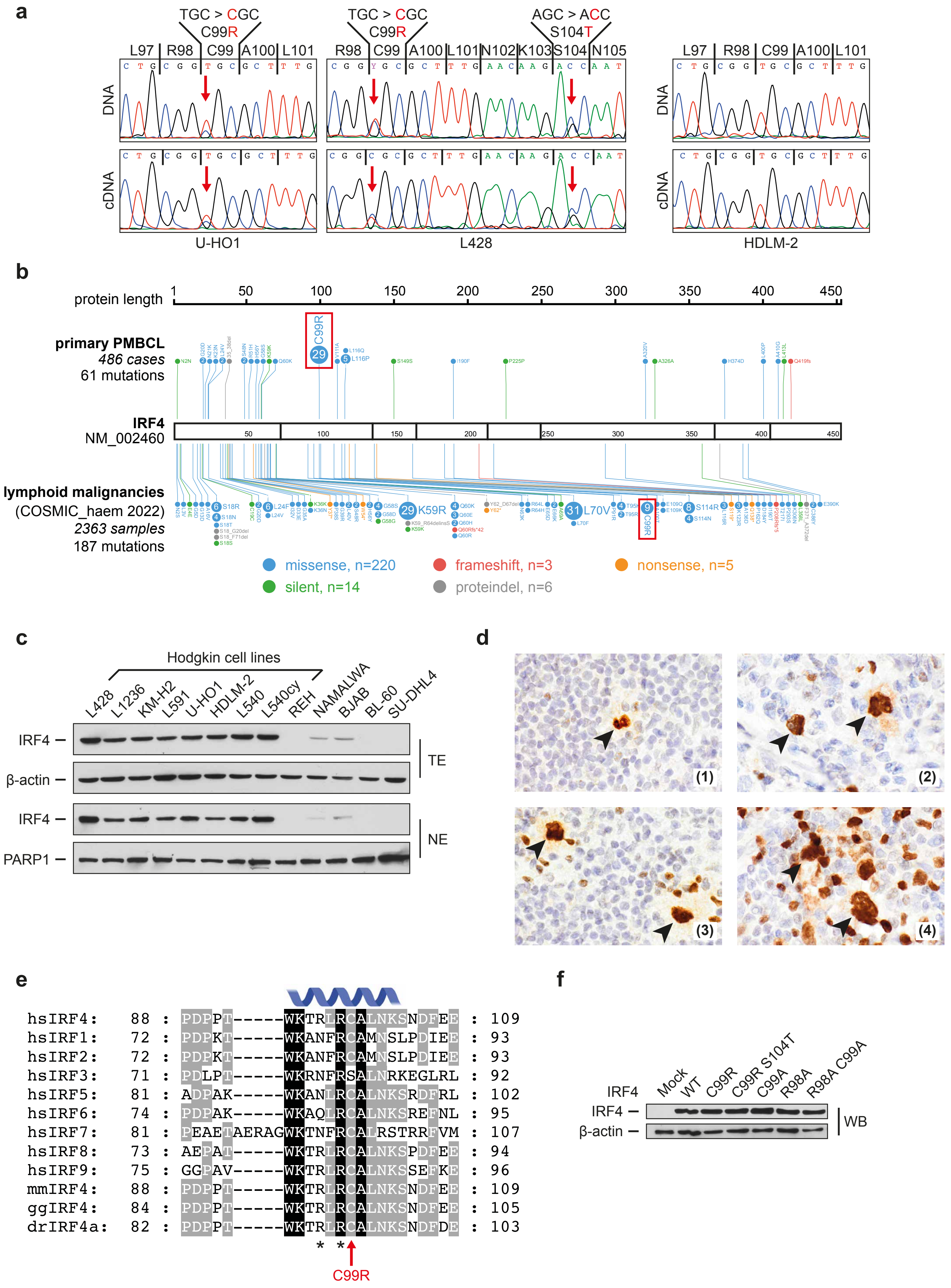

**Extended Data Fig. 1. IRF4 mutations and expression in human lymphoma.** (**a**) *IRF4* DNA (top panels) and cDNA (bottom panels) Sanger sequencing reads of the HRS cell lines U-HO1, harboring heterozygous IRF4-C99R, L428, harbouring heterozygous IRF4-C99R and S104T, and, as a control, HDLM-2 with IRF4-WT configuration. Red arrows indicate mutated nucleotide positions. (**b**) IRF4-mutation patterns in human lymphoma. Upper lollipop plots, IRF4-mutation frequencies in a cohort of 486 PMBCL. Lower lollipop plots, IRF4 mutation frequencies in human B cell lymphoma, numbers based on COSMIC database (https://cancer.sanger.ac.uk/cosmic; accessed on 2022/08/18) reporting 2363 samples tested. Of note, among the 9 samples harboring C99R reported in COSMIC, three samples are annotated as PMBCL and one as HL. (**c**) IRF4 immunoblotting of total extracts (TE, upper panels) or nuclear extracts (NE, lower panels) of various Hodgkin and non-Hodgkin cell lines, as indicated. Note, that HRS cell lines consistently show high IRF4 protein levels as compared to the non-Hodgkin cell lines. Data are representative of at least three independent experiments. (**d**) IRF4 immunohistochemistry of cHL. In total, all 90 analyzed cases (30 mixed cellularity, 30 nodular sclerosis, 30 lymphocyte-rich cHL cases) stained positive for IRF4. Representative IRF4 immunohistochemistries are shown: (1) nodular sclerosis cHL; (2) nodular sclerosis cHL; (3) lymphocyte-rich cHL; (4) mixed cellularity cHL. Magnification, 37,8X. (**e**) Protein alignment of human IRF1-9 as well as of mouse (Mus musculus; mm), chicken (Gallus gallus; gg) and zebrafish (Danio rerio; dr) IRF4 sequences corresponding to AA 88-109 of human IRF4. The position of C99 is indicated by a red arrow, two adjacent highly conserved arginines are marked by asterisks. The position of the corresponding IRF4 alpha3 DNA-recognition helix is shown in blue above the sequences. (**f**) Protein expression controls related to Fig. 1b. Nuclear extracts of HEK293 cells transfected with control plasmid (Mock) or the respective IRF4 variants, as indicated, were analyzed by immunoblotting for expression of IRF4 and, as a control, β-actin.

**Extended Data Figure 2**

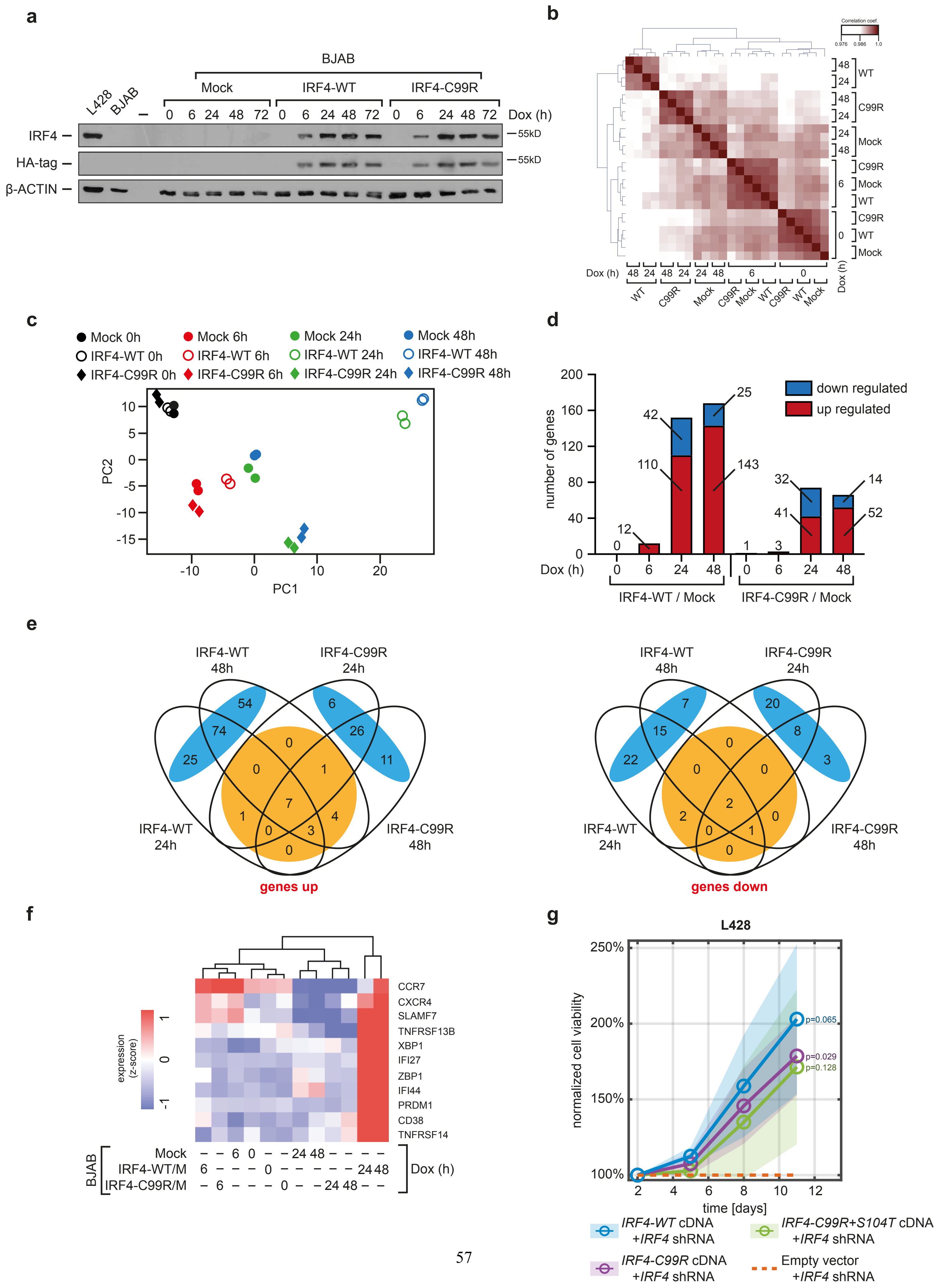

**Extended Data Fig. 2. IRF4-C99R functionality: IRF4-C99R regulates less and distinct genes compared to IRF4-WT and rescues HRS cells as efficiently as IRF4-WT following IRF4 knock-down.** (**a**) Mock, HA-tag-IRF4-WT and HA-tag-IRF4-C99R Tet-inducible non-Hodgkin BJAB cells were treated with doxycycline (Dox) for the indicated times. At each time, cells were analyzed by immunoblotting for IRF4 protein levels by use of antibodies to IRF4 or HA-tag. Extracts of untreated cHL L428 and non-Hodgkin BJAB cells (far left) as well as expression of β-actin were analyzed as positive or negative controls, respectively. One out of three independent experiments is shown. (**b**) Hierarchical clustering of Pearson correlation between the various transfectants, as indicated, across all expressed genes. Red colour indicates highly correlation between individual samples. Note, that control and respective IRF4 variants form separate clusters indicating differential gene regulation. (**c**) A two-dimension principal component plot between PC1 and PC2 for replicate samples across all expressed genes, again indicating differential gene expression between IRF4-WT and IRF4-C99R. (**d**) Following Dox-induction over time, bar diagrams show the number of differentially expressed genes that change expression two-fold between IRF4-WT *versus* Mock and IRF4-C99R *versus* Mock. Note, that IRF4-C99R regulates less genes up or down compared to IRF4-WT. (**e**) Four-way Venn diagram shows gene overlap between differentially expressed genes that change expression two-fold after 24 and 48 hours in IRF4-WT *versus* Mock and IRF4-C99R *versus* Mock cells. Left panel, up-regulated genes; right panel, down-regulated genes. Note, that for both classes of genes there is only marginal overlap between IRF4-WT- and IRF4-C99R-regulated genes (marked in orange), whereas both variants regulate individual gene sets (marked in blue). (**f**) Hierarchical clustering of standardized expression levels (z-score) of plasma cell genes. Red colours indicate highly expressed genes. Note, that IRF4-C99R is unable to induce plasma-cell genes, which are induced by IRF4-WT. (**g**) IRF4-WT and IRF4-C99R rescue HRS cells from *IRF4* shRNA-induced cell death. Following shRNA-mediated IRF4 knock-down, L428 HRS cells were transduced with empty vector (control), or cDNA encoding

IRF4-WT, IRF4-C99R, or IRF4-C99RS104T. Cell viability is shown normalized to control transduced cells. Note, that IRF4-C99R variants rescue the cells as efficiently as IRF4-WT from *IRF4* shRNA-induced cell death.

#### Extended Data Figure 3

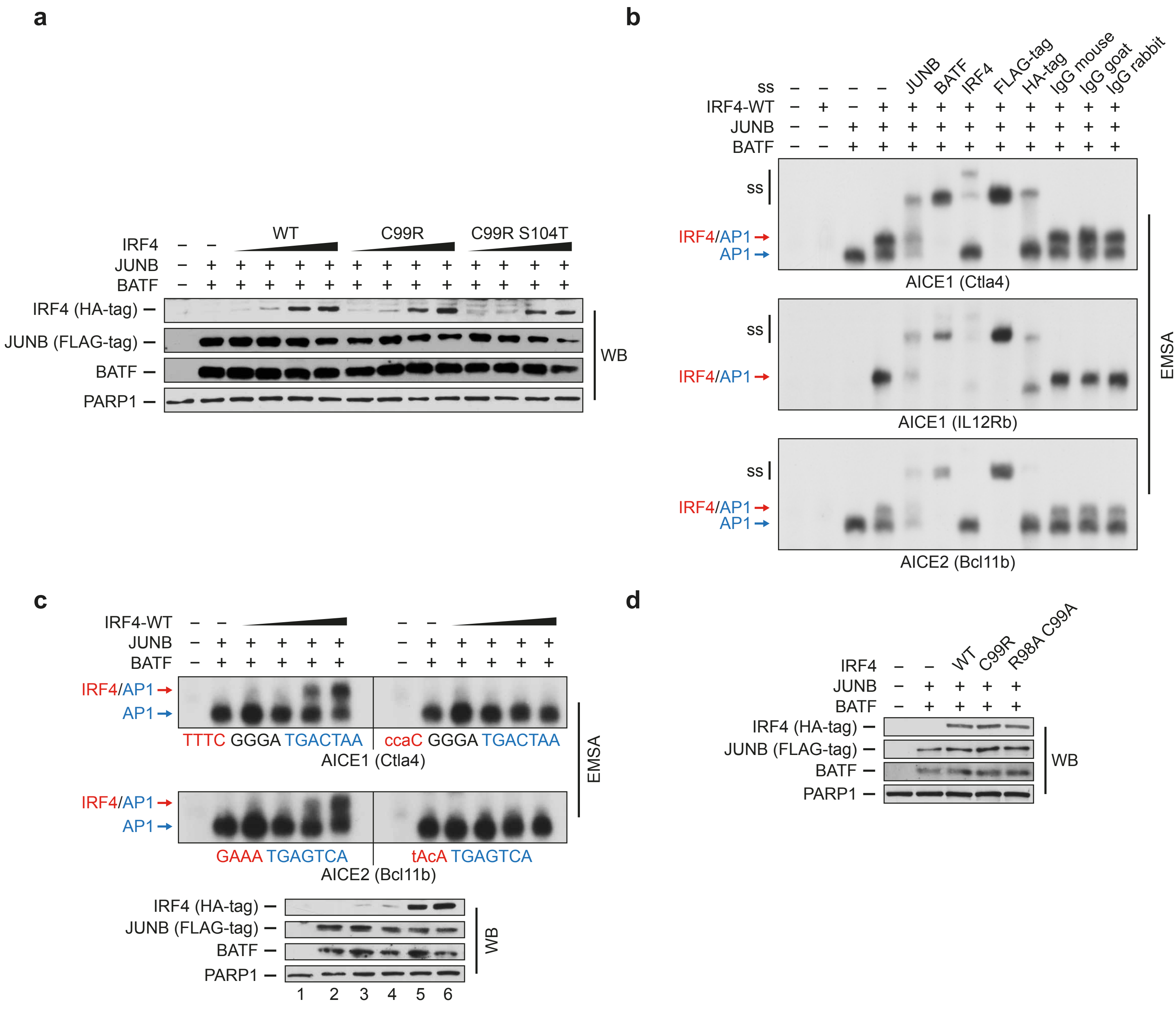

**Extended Data Fig. 3. Specificity of JUNB/BATF-IRF4 complexes binding to AICE motifs.** (**a**) Protein expression controls related to Fig. 1d. Nuclear extracts of HEK293 cells transfected with control plasmid (Mock; '-'), the respective IRF4 variants and JUNB and BATF were analyzed by immunoblotting for expression of IRF4, JUNB, BATF and, as a control, PARP1. (**b**) HEK293 cells were left untreated, or were transfected with IRF4-WT and/or JUNB and BATF, as indicated. Nuclear extracts were analyzed with or without addition of the indicated antibodies for supershift (ss) analyses at AICE1 (Ctla4) (top), AICE1 (IL12Rb) (center), and AICE2 (Bcl11b) (bottom). Red and blue arrows mark positions of IRF4-JUNB/BATF-DNA and JUNB/BATF-DNA complexes, respectively. (**c**) HEK293 cells were transfected with control plasmids (Mock, '-'), or were transfected with JUNB and BATF and increasing amounts of IRF4-WT. Nuclear extracts were analyzed by EMSA for binding at AICE1 (Ctla4) and AICE2 (Bcl11b) or variants thereof with mutated half-ISRE site. Note, that mutation of the respective ISRE binding motifs, indicated by small letters within the probe sequence, leads to complete loss of IRF4-JUNB-BATF composite complex formation at both sites (upper and lower right panels). In contrast, binding patterns of JUNB/BATF AP-1 complexes remain unchanged. Protein expression controls are shown underneath. Red and blue arrows mark positions of IRF4-JUNB/BATF-DNA and JUNB/BATF-DNA complexes, respectively. (**d**) Protein expression controls related to Fig. 1e. Nuclear extracts of HEK293 cells transfected with control plasmid (Mock) or the respective IRF4 variants and JUNB and BATF, as indicated, were analyzed by immunoblotting for expression of IRF4, JUNB, BATF and, as a control, PARP1.

#### Extended Data Figure 4

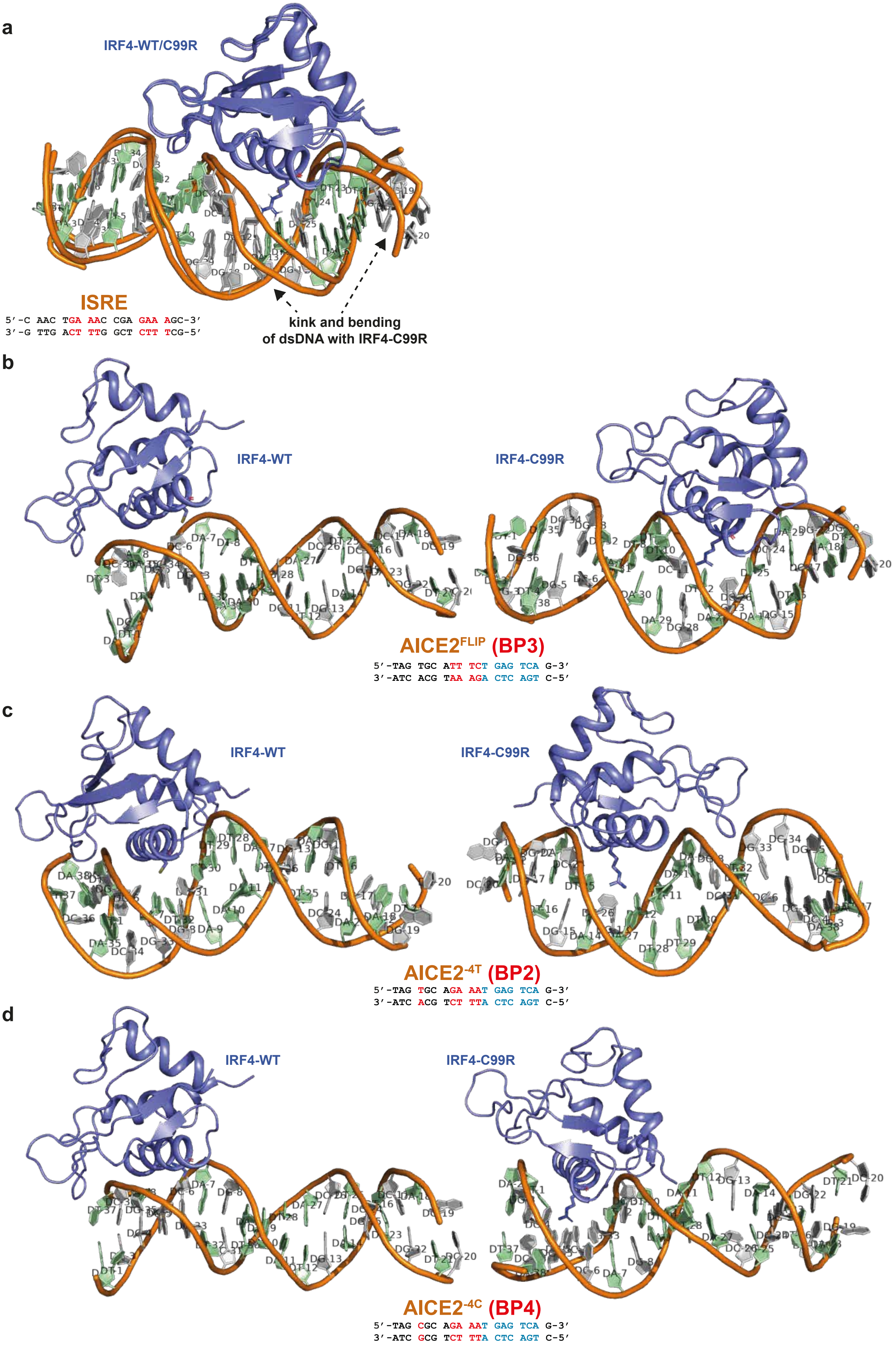

**Extended Data Fig. 4. Reference free DNA modelling and docking studies with IRF4-WT and IRF4-C99R at different DNA sequences.** (**a**) IRF4-C99R does not bind to intact ISRE DNA (see Fig. 1g) or needs to push the ISRE DNA for binding (shown here). The unbiased and reference free structural modelling of IRF4-C99R with the ISRE DNA shows bending of the DNA and displacement of the phosphate backbone to accommodate the recognition helix into the major groove of the DNA. Without this structural re-arrangement of ISRE DNA, binding of IRF4-C99R is difficult. (**b**) To understand the structural basis of the AICE2$^{FLIP}$ DNA interaction with IRF4-C99R but not IRF4-WT, we performed reference free DNA modeling and structural docking studies with IRF4-WT and IRF4-C99R. Our modelled and docked structures of AICE2$^{FLIP}$ DNA and IRF4-WT show no or minimal interactions despite the presence of major DNA grooves. These results are supported by scores/statistics of the docked models (Extended Data Table 4). Closer inspection of the mode of interaction showed that the DNA base pair stacking has moved upward in the structure. This may be one of the reasons why IRF4-WT could not bind to the AICE$^{FLIP}$ DNA as seen in the gel shift studies. In contrast, IRF4-C99R promotes bending of the AICE$^{FLIP}$ DNA enabling it to intercalate into the major grooves of DNA. We propose that the Arg residue facilitates minor changes to the DNA allowing engagement with the AICE$^{FLIP}$ DNA. (**c**, **d**) In order to elucidate the interactive properties of IRF4-WT and IRF4-C99R interactions with AICE2$^{-4T}$ (**c**) and AICE$^{-4C}$ (**d**) DNA elements, we performed reference free DNA modelling and docking studies. Foremost, we noted that the overall -4T and -4C DNA structures resembled AICE2 with distinct major and minor grooves. IRF4-C99R binds efficiently with both -4T and -4C DNA where the α-helix is well docked into the major groove of the DNAs with some differences in the DNA position and the orientation of IRF4, which reflected in the minor differences in the docking or interaction scores (c and d, right; Extended Data Table 4). Regarding IRF4-WT, it should be noted that the structural modeling and binding mode of IRF4-WT with the -4C DNA makes it difficult to confidently define a preferred binding mode. The best fit and docked structure shows the major

$\alpha$-helix of IRF4-WT is just stacked on the phosphate backbone and does not genuinely seem to be engaging in the interaction with the -4C DNA (d, left). To further explore the mode of binding, we analyzed all of the models in the designated clusters and re-calculated the unbiased reference free docking. These experiments confirmed that IRF4-WT has no preferred interaction mode with -4C DNA, which is stood out in the docking scores and statistics (Extended Data Table 4). This further corroborates our EMSA studies shown in Fig. 1F and assist in understanding why both variations in the DNA-fragment composition and mutations in IRF4 result in differential DNA-binding patterns. Blue/slate, IRF4 (both WT and C99R); orange, DNA phosphate backbone; pale green, dA and dT; grey, dG and dC.

#### Extended Data Figure 5

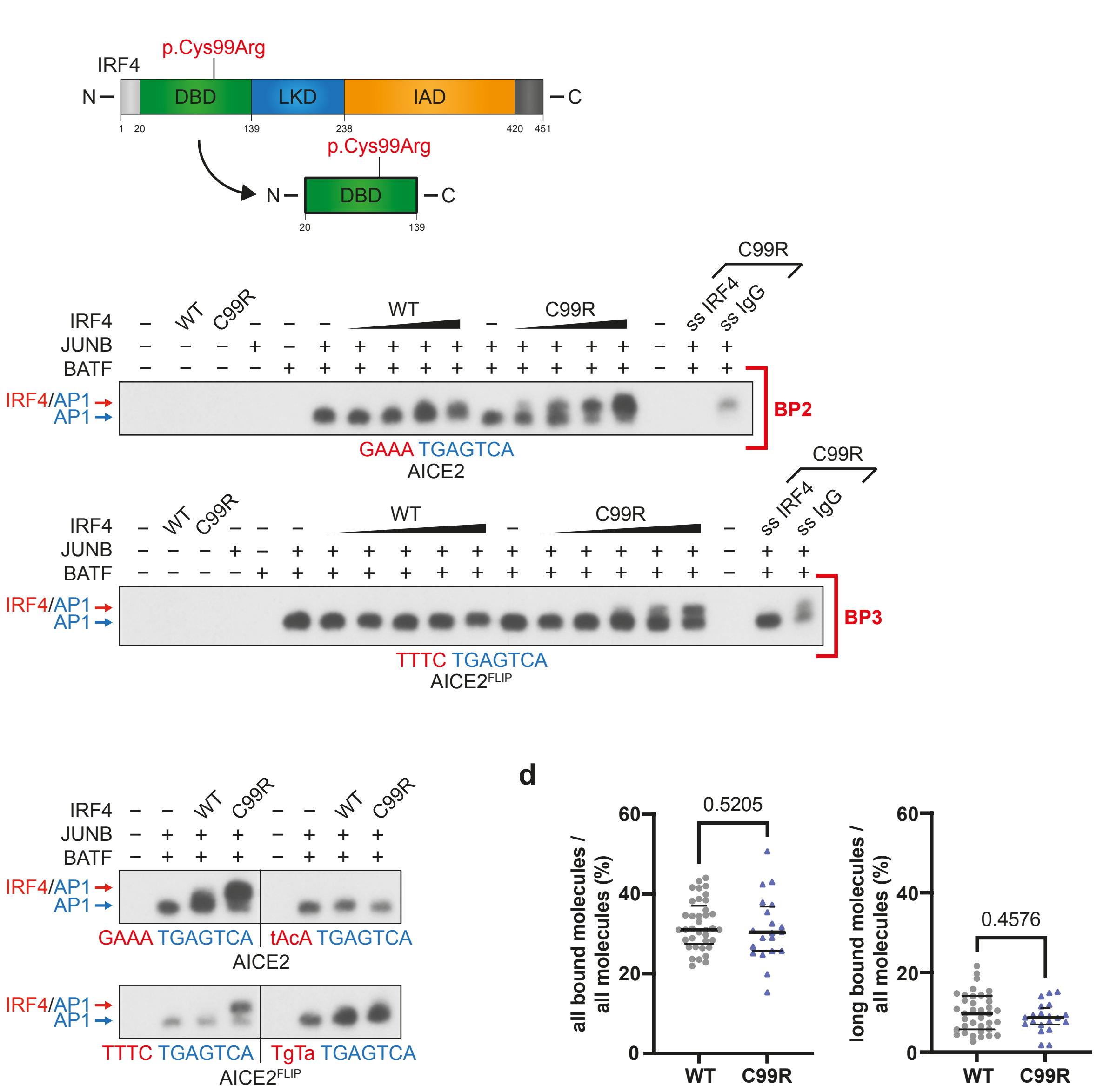

**Extended Data Fig. 5. IRF4 DNA binding studies by use of DNA binding domains only and by high-resolution microscopy.** (**a**) Schematic of full-length IRF4 (top) and IRF4-DBD (bottom), the latter used in Extended Data Figs. 5b and 5c. (**b**) Recombinant proteins comprising the DNA-binding domains (DBD) of JUNB (AA 269-329) or BATF (AA 28-87) were tested alone or together, or together in combination with increasing amounts of IRF4-WT or IRF4-C99R DBD (AA 20-139), as indicated, for their ability to bind to AICE2 (Bcl11b) (upper panel, BP2) or AICE2$^{FLIP}$ (lower panel, BP3). Red and blue arrows mark positions of IRF4-JUNB/BATF-DNA and JUNB/BATF-DNA complexes, respectively. Far right, supershift analyses by addition of antibody to IRF4 or, as a control, IgG. Note, that IRF4-C99R DBD binds, compared to IRF4-WT, much stronger to AICE2 (Bcl11b), and exclusively to AICE2$^{FLIP}$. (**c**) Recombinant proteins, as described in (b) were analyzed for binding to AICE2 (Bcl11b)-WT (upper left) and AICE2 (Bcl11b)-IRFMut (upper right), as well as to AICE2$^{FLIP}$ (lower left) and AICE2$^{FLIP}$-IRFMut (lower right). Note, that mutation of the IRF motif in each of the probes abolishes formation of IRF4-JUNB/BATF-DNA composite complexes. Positions of the complexes are indicated as described in (b). (**d**) Determination of IRF4-WT and -C99R binding fractions. Fractions of all bound SiR-HaloTag-IRF4-WT and IRF4–C99R molecules (left) and molecules bound for > 2s (right) as determined by single-molecule fluorescence microscopy with interlaced time-lapse illumination.

#### Extended Data Figure 6

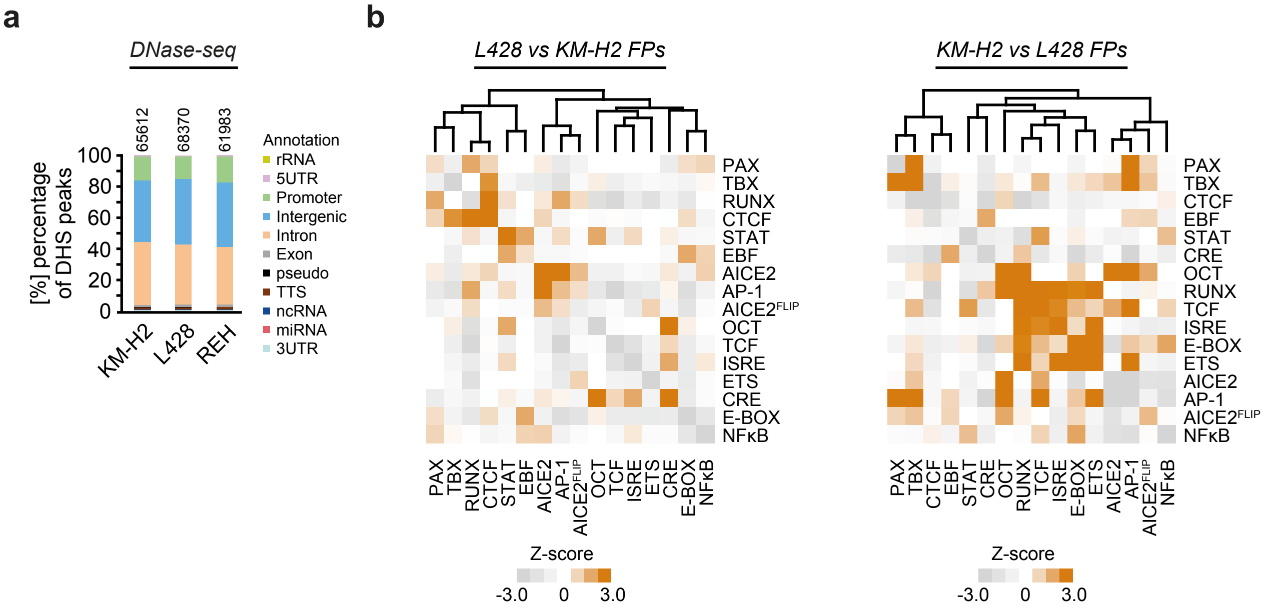

**Extended Data Fig. 6. Genome-wide localization of DNaseI-Seq peaks and co-occurring clustering of AICE2.** (**a**) Genome-wide localization of DNase-seq peaks in KM-H2$^{IRF4\text{-}WT}$, L428$^{IRF4\text{-}C99R}$ and REH cells. Note, that most peaks are intergenic or intronic, with about 20% of DHSs showing 15-20% promoter enrichment. (**b**) Heatmap detailing z-scores of self- and co-occurrence enrichments in L428 *versus* KM-H2 (left) or KM-H2 *versus* L428 (right) footprints for 16 motifs corresponding to TFs important to B cell and HL gene regulation. |Z-scores| above 1.96 represent ≥2σ enrichment.

#### Extended Data Figure 7

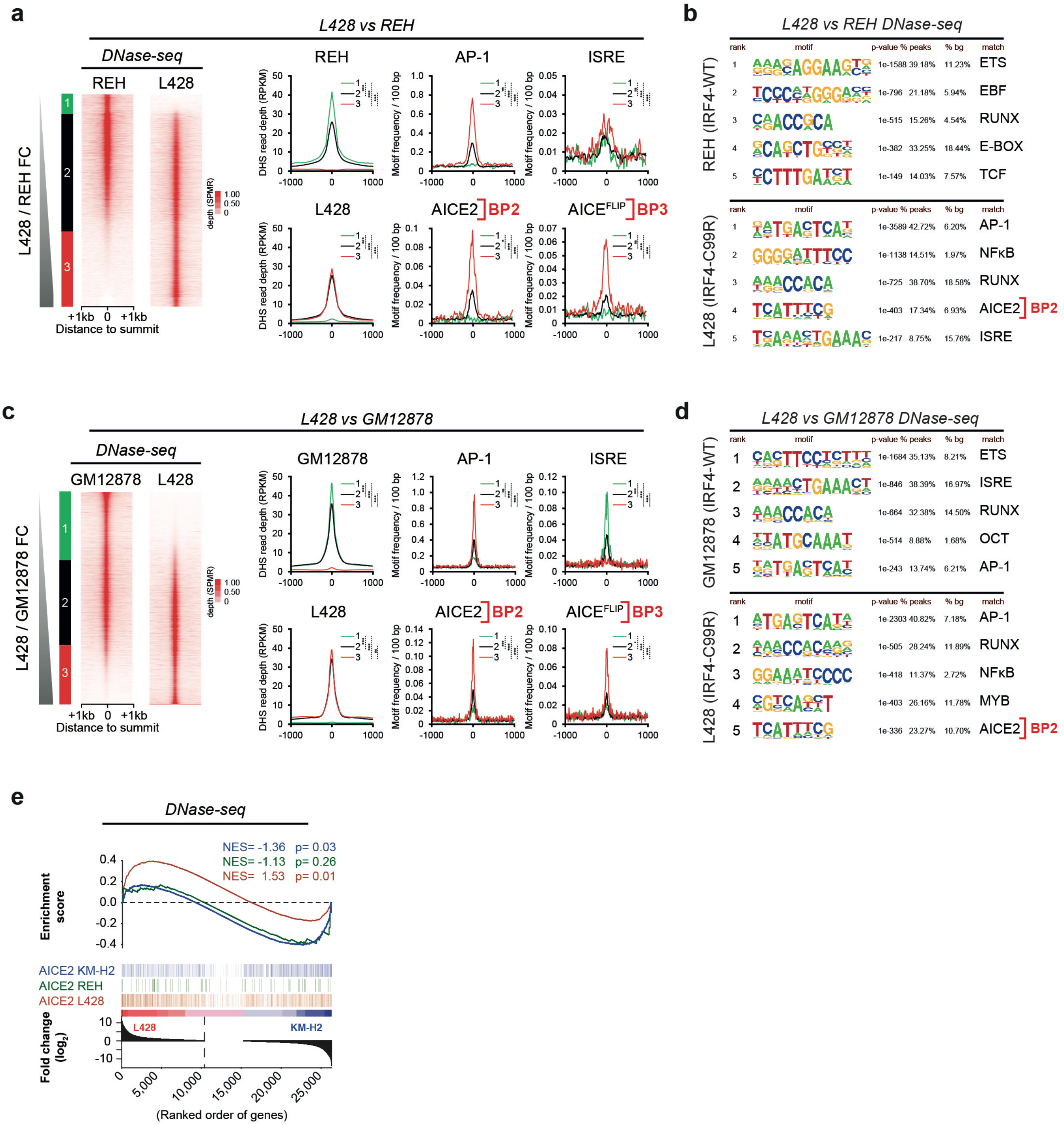

**Extended Data Fig. 7. AICE2 motifs are enriched in IRF4-C99R HL cells and are associated with increased gene expression.** (**a**) Heatmaps showing the DNase-seq fold change analysis (left) between L428$^{\text{IRF4-C99R}}$ HL and non-Hodgkin REH cells and corresponding DNase-Seq motif average profiles (right). (**b**) HOMER *de novo* motif discovery-results from classes 1 (REH-specific, top) and 3 (L428$^{\text{IRF4-C99R}}$-specific, bottom), defined in (a). (**c**, **d**) Same as (a, b) but comparing L428 with GM12878 DNase-Seq. Note, that both comparisons reveal AICE2 and AICE2$^{\text{FLIP}}$ motifs as specifically enriched in L428 $^{\text{IRF4-C99R}}$ cells. (**e**) GSEA of footprinted L428$^{\text{IRF4-C99R}}$, KM-H2$^{\text{IRF4-WT}}$, and REH AICE2 motif presence (left, center, right) versus L428 / KM-H2 RNA-seq fold change. Note, that footprinted AICE2 motif presence is associated with increased gene expression for L428 followed by KM-H2, but not in REH cells.

#### Extended Data Figure 8

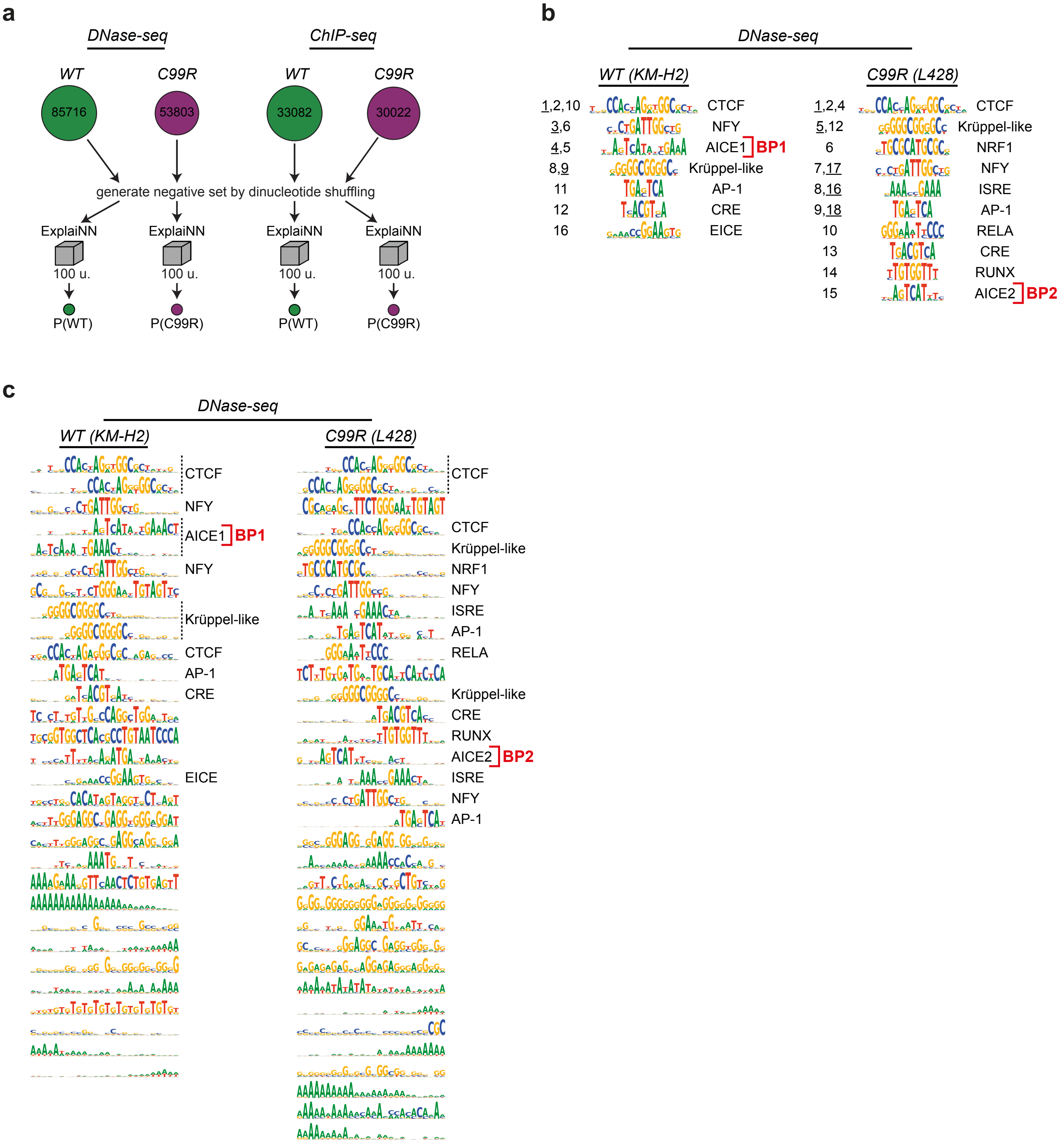

**Extended Data Fig. 8.** (**a**) The four ExplaiNN models trained in this work. (**b**) *De novo* motif discovery in specific DHSs datasets of KM-H2$^{IRF\text{-}WT}$ and L428$^{IRF4\text{-}C99R}$ using ExplaiNN. Motifs are ranked by their importance (left). When more than one motif of the same class was identified, the rank of the displayed motif is underlined. Only motifs that could by annotated with a biological representation are shown. (**c**) List if all motifs identified in the DHS data sets by ExplaiNN in (b) ranked by their importance.

#### Extended Data Figure 9

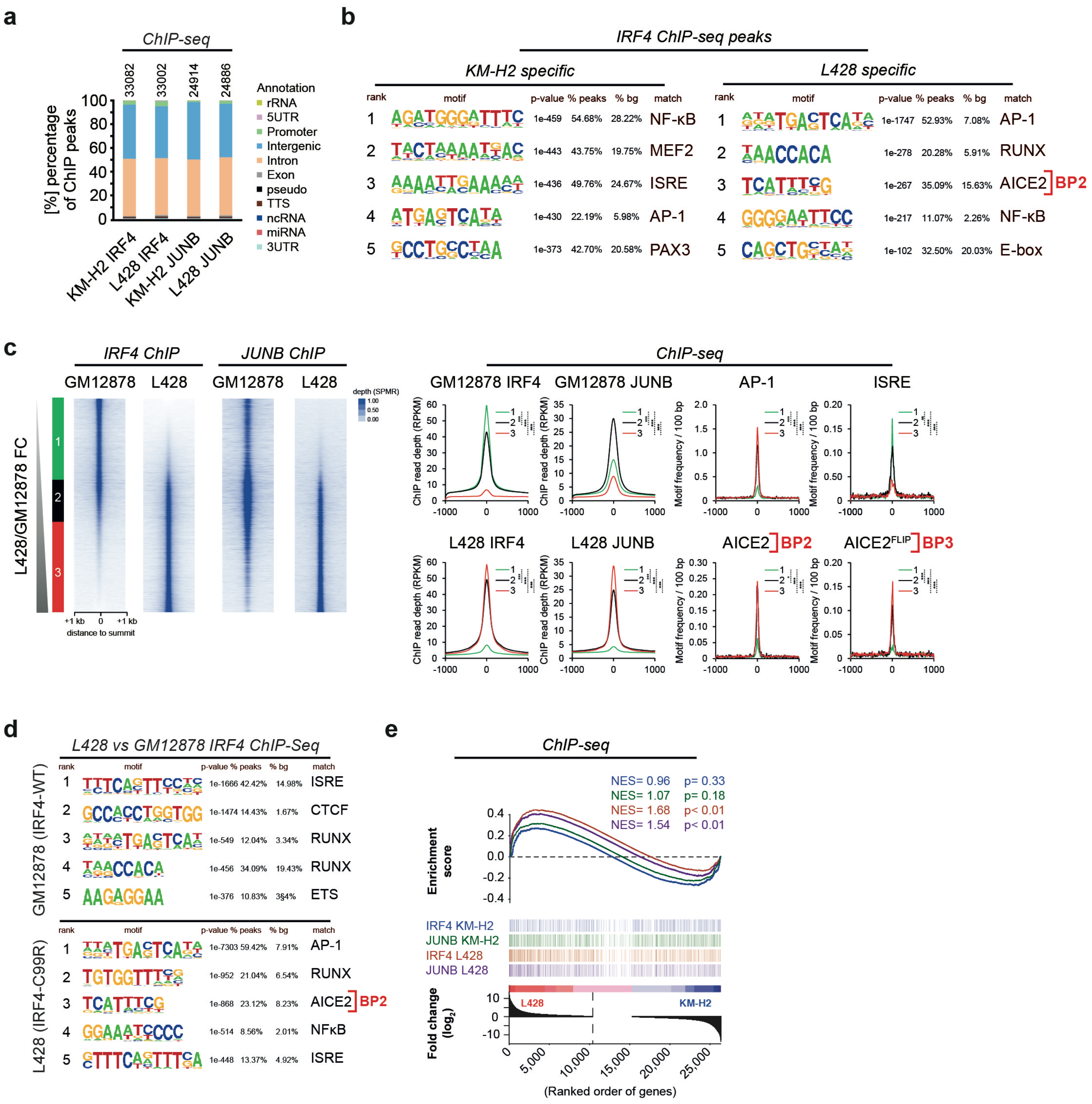

**Extended Data Fig. 9. AICE2 motifs are associated with increased IRF4/JUNB co-localization in IRF4-C99R HL cells.** (**a**) Genome-wide localization of IRF4, JUNB ChIP-Seq peaks in KM-H2 and L428 cells. Most peaks are intergenic and intronic. (**b**) HOMER motif discovery results in KM-H2 (left)- and L428 (right)-specific IRF4 ChIP peaks defined in Fig. 2g. (**c**) Heatmaps showing IRF4 ChIP fold change (FC) analysis (blue, leftmost) between L428 and GM12878 cells, corresponding JUNB ChIP-Seq signal (blue, rightmost) and corresponding ChIP-seq motif average profiles (right). Note, that AP-1, AICE2 (BP2) and AICE$^{FLIP}$ (BP3) signals are highest in L428$^{IRF4\text{-}C99R}$-specific peaks. (**d**) HOMER *de novo* motif discovery-results from classes 1 and 3 (top and bottom, respectively) in GM12878 (top) and L428 (bottom)-specific IRF4 ChIP-seq peaks defined in (c). (**e**) GSEA of the top 1,000 differential IRF4 and JUNB L428 and KM-H2 ChIP-seq peaks (left and right, respectively) against corresponding L428 / KM-H2 RNA-seq fold change (FC). Note, that IRF4 and JUNB ChIP peaks are only significantly associated with increased gene expression in L428$^{IRF4\text{-}C99R}$ cells. Significance levels: *, p<0.05; **, p<0.01; ***, p<0.001; ns, non-significant.

### Extended Data Figure 10

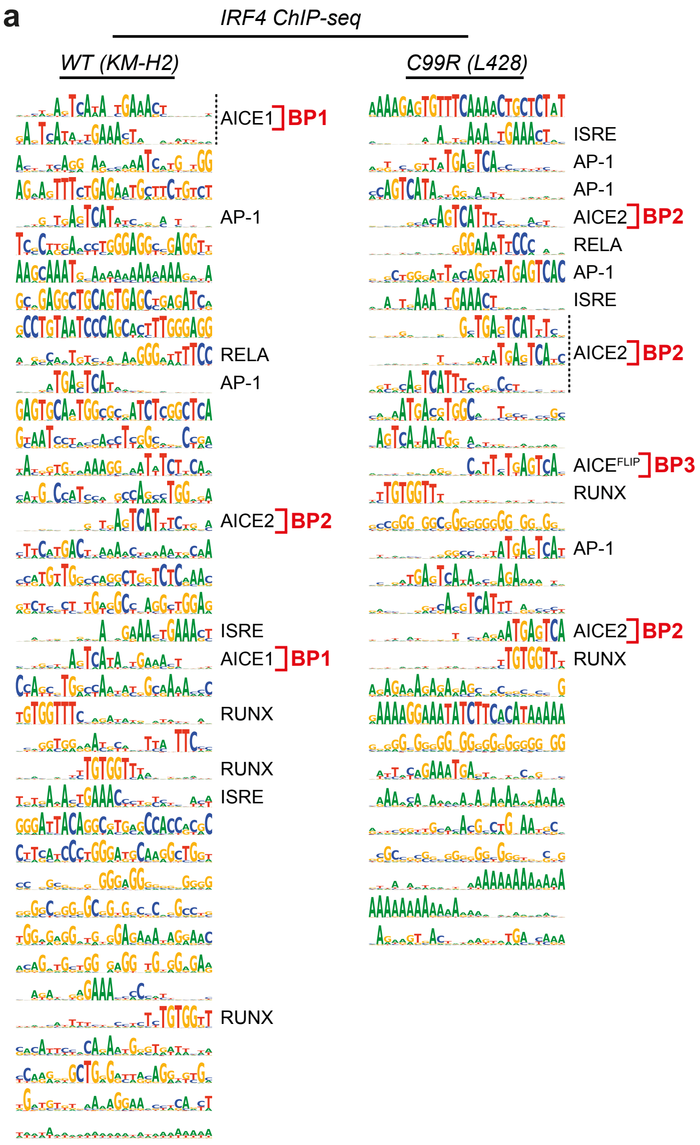

**Extended Data Fig. 10. Increased AICE2 motif frequencies in IRF4-C99R HL cells from neural-network based motif discovery approaches.** (**a**) Full report of motif discovery results of the ChIP-seq data sets using ExplaiNN, detailing sub-motifs making aggregate motifs described in Figure 2i. Note, that AICE1 (BP1) is only identified in KM-H2$^{IRF4\text{-}WT}$ cells, whereas AICE2 BP2 is overrepresented and AICE2 BP3 exclusively found in L428$^{IRF\text{-}C99R}$ cells.

#### Extended Data Figure 11

a

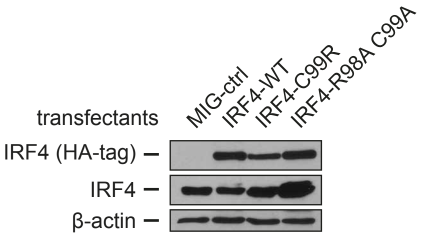

b

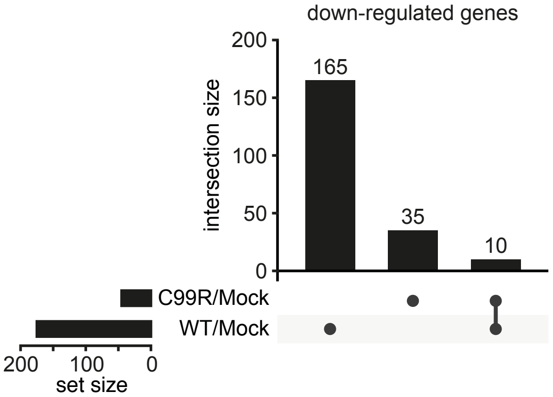

**Extended Data Fig. 11. Expression controls of transduced C57BL/6 mouse splenic B cells and IRF4-C99R down-regulated genes.** (**a**) C57BL/6 mouse splenic B cells transduced with MIG control retrovirus (MIG-ctrl; Mock), IRF4-WT, IRF4-C99R or, as a further control, IRF4-R98AC99A, were analyzed by immunoblotting for expression levels of IRF4 by use of HA-tag and IRF4 antibody, respectively. β-actin was analyzed as a control. (**b**) IRF4-C99R and IRF-WT regulated genes show only low overlap, as shown in UpSet plots for down-regulated genes.

#### Extended Data Figure 12

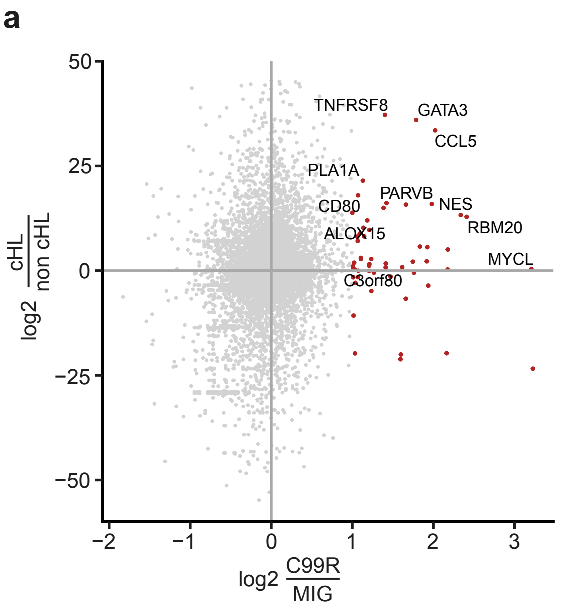

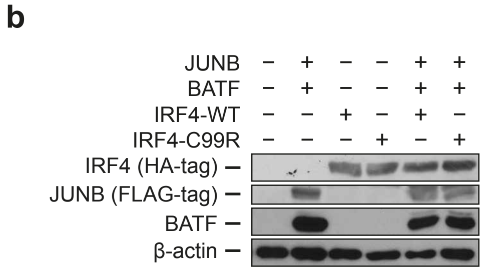

**Extended Data Fig. 12. IRF4-C99R up-regulated genes encompass cHL hallmark genes. Expression controls for reporter gene studies.** (**a**) Comparison of fold changes between IRF4-C99R-induced genes with differentially expressed genes of Hodgkin and non-Hodgkin cell lines based on RNA-seq analyses. (**b**) Protein expression controls related to Fig. 4d. Whole cell extracts of HEK293 cells transfected with control plasmid (Mock), JUNB, BATF, IRF4-WT or IRF4-C99R alone or in combination, as indicated, were analyzed by immunoblotting for expression of JUNB, BATF, IRF4 and, as a control, β-actin.

**Extended Data Table 1. Features of the IRF4 mutation.**

| Variant Annotation | |
|---|---|
| **Chromosome** | 6 |
| **Genomic Position (GRCh38)** | 394899 |
| **cDNA Position (NM_001195286)** | 295 |
| **Nucleotide Reference** | T |
| **Nucleotide variant** | C |
| **Protein Variant** | Cys99Arg; C99R |
| **dbSNP153** | No entry |
| **gnomAD** | No entry |
| **COSMIC (v96)** | Somatic report, 9 entries (Mutation ID COSV66704715) |

| *In silico* Pathogenicity Prediction Models | |
|---|---|
| **CADD** | 24.0 |
| **SIFT** | Deleterious (0) |
| **Polyphen-2** | Probably damaging (1) |
| **LRT** | Deleterious (0) |
| **MutationTaster** | Disease causing (1) |
| **PROVEAN** | Deleterious (-11.56) |
| **MetaSVM** | Deleterious (1.011) |
| **M-Cap** | Possibly pathogenic (0.87) |
| **fathmm_MKL-coding** | Deleterious (0.942) |

**Extended Data Table 2. *IRF4* exon 3 mutation analysis of HRS cells.**

| sample no | disease stage | subtype | EBV | mutated sequences per total sequences* | c.295T>C |
|---|---|---|---|---|---|
| **1** | primary | NS | - | 0 / 4 | wt |
| **2** | primary | NS | - | 0 / 2 | wt |
| **3** | primary | MC | - | 0 / 2 | wt |
| **4** | primary | NS | - | 0 / 2 | wt |
| **5** | primary | NS | - | 0 / 2 | wt |
| **6** | primary | MC | + | 0 / 2 | wt |
| **7** | primary | NS | - | 0 / 4 | wt |
| **8** | primary | NS | - | 0 / 2 | wt |
| **9** | primary | NS | - | 0 / 3 | wt |
| **10** | primary | NS | - | 0 / 2 | wt |
| **11a** | primary | NS | - | 2 / 4 | mut |
| **11b** | relapse | NS | - | 2 / 3 | mut** |
| **12** | relapse | NS | - | 0 / 4 | wt |
| **13** | relapse | NS | - | 0 / 2 | wt |
| **14** | relapse | NS | - | 0 / 3 | wt |
| **15** | relapse | NS | - | 0 / 4 | wt |
| **16** | relapse | NS | - | 2 / 3 | mut |
| **17** | relapse | MC | + | 0 / 3 | wt |
| **18** | relapse | NS | - | 2 / 5 | mut |
| **19** | relapse | LR*** | + | 0 / 3 | wt |
| **20** | relapse | MC | + | 0 / 2 | wt |

NS, nodular sclerosis; MC, mixed cellularity; LR, lymphocyte-rich; wt, wild type; mut, mutated.

* 2-4 informative sequences from pools of 10 microdissected HRS cells were analyzed per case.

**PCR products were cloned into pGEMTeasy and sequenced from plasmid DNA to identify 4/27 mutated sequences.

***subtype at primary diagnosis

**Extended Data Table 3. (separate file)**
Differentially expressed genes in IRF4-WT and IRF4-C99R BJAB cells.

**Extended Data Table S4. Validation report for the predicted structure of WT/C99R IRF4 with different DNA fragments.** The table shows the docking results with indicated scores reflecting the best or top complex structure cluster. The tabulated scores are primarily derived from docking scores (HADDOCK) while additional scores denote RMSD, electro-static potential, van der Waals or buried interphase and the z-scores of the predicted complexes. The Z-score indicates the cluster's HADDOCK score from the average of all clusters. A lower Z-score denotes a superior model.

| | **ISRE-C99R** | **AICE2$^{-4T}$-WT** | **AICE2$^{-4T}$-C99R** | **AICE2$^{-4C}$-WT** | **AICE2$^{-4C}$-C99R** | **AICE2$^{FLIP}$-WT** | **AICE2$^{FLIP}$-C99R** | **AICE1-C99R** | **AICE2-C99R** |
|---|---|---|---|---|---|---|---|---|---|
| **HADDOCK score** | -163.2 +/- 9.7 | -99.0 +/- 3.2 | -105.4 +/- 11.1 | -68.9 +/- 10.4 | -119.1 +/- 12.1 | -78.7 +/- 7.4 | -94.5 +/- 2.5 | -95.6 +/- 0.2 | -129.1 +/- 12.1 |
| **Cluster size** | 28 | 11 | 15 | 14 | 13 | 9 | 16 | 12 | 15 |
| **RMSD** | 1.0 +/- 0.6 | 9.0 +/- 0.2 | 10.2 +/- 0.2 | 6.2 +/- 0.3 | 0.8 +/- 1.2 | 13.8 +/- 0.4 | 9.9 +/- 0.1 | 18.2 +/- 0.0 | 0.8 +/- 1.2 |
| **Van der Waals** | -86.5 +/- 8.0 | -57.2 +/- 8.0 | -59.8 +/- 6.3 | -48.4 +/- 9.1 | -60.8 +/- 7.0 | -46.2 +/- 8.4 | -54.3 +/- 2.9 | -46.5 +/- 1.1 | -60.8 +/- 7.0 |
| **Electrostatic Energy** | -528.0 +/- 38.7 | -254.2 +/- 14.7 | -288.4 +/- 19.5 | -175.0 +/- 11.3 | -382.2 +/- 41.0 | -197.3 +/- 42.6 | -218.6 +/- 10.2 | -293.2 +/- 7.4 | -382.2 +/- 41.0 |
| **Desolvation Energy** | 25.0 +/- 2.8 | 8.8 +/- 3.4 | 7.6 +/- 1.6 | 4.3 +/- 2.5 | 17.9 +/- 3.6 | -0.2 +/- 2.2 | 3.3 +/- 2.3 | 8.0 +/- 0.7 | 17.9 +/- 3.6 |
| **Restraints violation** | 38.8 +/- 24.3 | 2.3 +/- 0.1 | 44.6 +/- 14.7 | 2.2 +/- 0.3 | 2.9 +/- 0.1 | 11.1 +/- 8.7 | 2.1 +/- 1.1 | 15.7 +/- 16.4 | 2.9 +/- 0.1 |
| **Buried Surface Area** | 2315.8 +/- 170.7 | 1342.1 +/- 67.0 | 1415.1 +/- 74.9 | 1210.0 +/- 114.6 | 1695.7 +/- 217.4 | 982.2 +/- 152.9 | 1299.5 +/- 117.0 | 1094.0 +/- 19.5 | 1695.7 +/- 217.4 |
| **Z-Score** | -1.8 | -1.5 | -0.8 | -0.2 | -2.3 | -0.3 | -1.4 | +0.2 | -2.1 |

**Extended Data Table 5. (separate file)**
Differentially expressed genes in C57BL/6 splenic B cells transduced with IRF4 variants.

**Extended Data Table 6. Oligonucleotides used in the present study.**

| Probes for EMSA | | |
|---|---|---|
| ISRE (ISG15) | forward | AGCTGGGAAAGGGAAACCGAAACTG |
| | reverse | AGCTCAGTTTCGGTTTCCCTTTCCC |
| AICE1 (Ctla 4) | forward | AGCTCTTGCCTTAGAGGTTTCGGGATGACTAATACT GTA |
| | reverse | AGCTTACAGTATTAGTCATCCCGAAACCTCTAAGGC AAG |
| AICE1 (Il12Rb) | forward | AGCTGCTTTTGCTTTCACTTTGACTGGCCTGGAGAC AATGAGTT |
| | reverse | AGCTAACTCATTGTCTCCAGGCCAGTCAAAGTGAAA GCAAAAGC |
| AICE2 (Bcl11b) | forward | AGCTTAGTGCAGAAATGAGTCAGAGATCAAAGAAG |
| | reverse | AGCTCTTCTTTGATCTCTGACTCATTTCTGCACTA |
| SP1 | forward | AGCTATTCGATCGGGGCGGGGCGAGC |
| | reverse | AGCTGCTCGCCCCGCCCCGATCGAAT |
| AICE2 5´TTTC | forward | AGCTTAGTGCATTTCTGAGTCAGAGATCAAAGAAG |
| | reverse | AGCTCTTCTTTGATCTCTGACTCAGAAATGCACTA |
| AICE2 3´GAAA | forward | AGCTTAGTGCATGAGTCAGAAAGAGATCAAAGAAG |
| | reverse | AGCTCTTCTTTGATCTCTTTCTGACTCATGCACTA |
| AICE2 3´TTTC | forward | AGCTTAGTGCATGAGTCATTTCGAGATCAAAGAAG |
| | reverse | AGCTCTTCTTTGATCTCGAAATGACTCATGCACTA |
| AICE2-4C (Bcl11b) | forward | AGCTTAGCGCAGAAATGAGTCAGAGATCAAAGAAG |
| | reverse | AGCTCTTCTTTGATCTCTGACTCATTTCTGCGCTA |
| AICE2 (Bcl11b_short) | forward | AGTGCAtAcATGAGTCAGAGATC |
| | reverse | AGGATCTCTGACTCATgTaTGCA |
| AICE2 (Bcl11b_short_IRFm ut) | forward | AGTGCAGccATGAGTCAGAGATC |
| | reverse | CGGATCTCTGACTCATggCTGCA |
| AICE$^{FLIP}$ (Bcl11b_5´TTTC_F LIP_short) | forward | AGTGCATTTCTGAGTCAGAGATC |
| | reverse | AGGATCTCTGACTCAGAAATGCA |
| | forward | AGTGCATgTaTGAGTCAGAGATC |

| | | |
|---|---|---|
| AICE$^{FLIP}$ (Bcl11b_5´TTTC_FLIP_short_IRFmut) | reverse | AGGATCTCTGACTCAtAcATGCA |
| GATA3Peak_1 | forward | AGCTACTGAATGAGTCAGAGAGGCCCAAA |
| | reverse | AGCTTTTGGGCCTCTCTGACTCATTCAGT |
| GATA3Peak_2 | forward | AGCTAGTCATAAATGAGTCATGGC |
| | reverse | AGCTGCCATGACTCATTTATGACT |
| GATA3Peak_3 | forward | AGCTAATGCAGGAATGACTCACTTG |
| | reverse | AGCTCAAGTGAGTCATTCCTGCATT |
| Primers for Site-Directed Mutagenesis | | |
| IRF4mut_C99R | forward | GAAGACGCGCCTGCGGCGCGCTTTGAACAAGAGCAATG |
| | reverse | CATTGCTCTTGTTCAAAGCGCGCCGCAGGCGCGTCTTC |
| IRF4mut_C99RS104T | forward | CGCCTGCGGCGCGCTTTGAACAAGACCAATGACTTTG |
| | reverse | CAAAGTCATTGGTCTTGTTCAAAGCGCGCCGCAGGCG |
| IRF4mut_C99A | forward | GGAAGACGCGCCTGCGGGCCGCTTTGAACAAGAG |
| | reverse | CTCTTGTTCAAAGCGGCCCGCAGGCGCGTCTTCC |
| IRF4mut_R98A | forward | GAAGACGCGCCTGGCGTGCGCTTTGAACAAGAG |
| | reverse | CTCTTGTTCAAAGCGCACGCCAGGCGCGTCTTC |
| IRF4mut_R98AC99A | forward | GGAAGACGCGCCTGGCGGCCGCTTTGAACAAGAGCAATG |
| | reverse | CATTGCTCTTGTTCAAAGCGGCCGCCAGGCGCGTCTTCC |

**Extended Data Table 7. Antibodies used in the present study.**

| ANTIBODIES | COMPANY | IDENTIFIER |
|---|---|---|
| Antibodies for EMSA | | |
| IRF4 (M-17) | Santa Cruz Biotechnology | sc-6059 |
| JUNB (N-17) | Santa Cruz Biotechnology | sc-46 |
| BATF (WW-8) | Santa Cruz Biotechnology | sc-100974 |
| HA-tag | Cell Signaling Technology | 3724 |
| Anti-Flag M2 | Sigma-Aldrich | F1804 |
| BATF (D7C5) | Cell Signaling Technology | 8638 |
| IgG1 mouse | R&D Systems | MAB002 |
| IgG rabbit | R&D Systems | AB-105-c |
| IgG goat | R&D Systems | AB-108-c |
| Antibodies for Western Blot | | |
| IRF4 (M-17) | Santa Cruz Biotechnology | sc-6059 |
| IRF4 (D9P5H) | Cell Signaling Technology | #15106 |
| HA-probe | Santa Cruz Biotechnology | sc-805 |
| HA-tag | Cell Signaling Technology | #3724 |
| Anti-Flag M2 | Sigma-Aldrich | F1804 |
| BATF (D7C5) | Cell Signaling Technology | #8638 |
| PARP1 | Santa Cruz Biotechnology | sc-8007 |
| β-actin | Sigma-Aldrich | #A5316 |
| Antibodies for FACS | | |
| CD138-PE | Biolegend | 142504 |
| PerCP/cy5.5 Anti-mouse B220 | Biolegend | 103235 |
